# Annual dynamics of *Zymoseptoria tritici* populations in wheat cultivar mixtures: a compromise between the efficiency and durability of a recently broken-down resistance gene?

**DOI:** 10.1101/2021.04.23.441180

**Authors:** Carolina Orellana-Torrejon, Tiphaine Vidal, Anne-Lise Boixel, Sandrine Gélisse, Sébastien Saint-Jean, Suffert Frédéric

## Abstract

Cultivar mixtures slow polycyclic epidemics but may also affect the evolution of pathogen populations by diversifying the selection pressures exerted by their plant hosts at field scale. We compared the dynamics of natural populations of the fungal pathogen *Zymoseptoria tritici* in pure stands and in three binary mixtures of wheat cultivars (one susceptible cultivar and one cultivar carrying the recently broken-down *Stb16q* gene) over two annual field epidemics. We combined analyses of population ‘size’ based on disease severity, and of population ‘composition’ based on changes in the frequency of virulence against *Stb16q* in seedling assays with more than 3000 strains. Disease reductions were observed in mixtures late in the epidemic, at the whole canopy scale and on both cultivars, suggesting the existence of a reciprocal ‘protective’ effect. The three cultivar proportions in the mixtures (0.25, 0.5 and 0.75) modulated the decrease in (i) the size of the pathogen population relative to the two pure stands, (ii) the size of the virulent subpopulation, and (iii) the frequency of virulence relative to the pure stand of the cultivar carrying *Stb16q*. Our findings suggest that optimal proportions may differ slightly between the three indicators considered. We argued potential trade-offs that should be taken into account when deploying a resistance gene in cultivar mixtures: between the dual objectives ‘efficacy’ and ‘durability’, and between the ‘size’ and ‘frequency’ of the virulent subpopulation. Based on current knowledge, it remains unclear whether virulent subpopulation size or frequency has the largest influence on interepidemic virulence transmission.

## Introduction

The industrialisation of agrosystems has greatly simplified the implementation of crop protection. In conventional cereal farming, fungal disease control is based on two major pillars: phytosanitary products and disease-tolerant or resistant cultivars. Most fields are planted with a single cultivar, in which effective resistance usually depends on the combination of a few qualitative resistance genes rather than diversified quantitative genetic components. The strong resistance conferred by a single gene is simpler for breeders to handle, but may be easily overcome by pathogens (McDonald & Linde, 2002).

The number of wheat cultivars grown in North-Western Europe is small (Perronne *et al*., 2017). This leads to the exertion of unidirectional selection pressures on plant pathogen populations, fostering the breakdown of qualitative resistances, and resulting in an increase in the frequency of virulent strains just after their emergence (Brown & Tellier, 2011). Host homogeneity thus limits the ‘durability’ of qualitative resistance, defined as its efficacy over a prolonged period of widespread use under conditions conducive to the disease (Johnson, 1984). Pathogen evolution, which occurs at a variable rate, usually results in the replacement of the ‘oldest’ cultivars grown with others carrying ‘new’ sources of resistance. These dynamics have been widely described in publications on wheat diseases as a response to the common ‘boom-and-bust’ cycles observed in clonal pathogen populations (e.g. McIntosh & Brown, 1997) and to the more gradual breakdown of resistance in diversified sexual populations (e.g. Cowger *et al*., 2000). In this context, one of the most striking issues currently facing us is the need to reconcile the efficiency and durability of the resistances available to farmers, by identifying optimal strategies for their deployment over different space-time scales (Fabre *et al*., 2015).

Increasing crop diversity at field scale has been widely reported to result in more efficient control over plant disease epidemics. This is the case for cultivar mixtures (Wolfe, 1985), in which cultivars of the same species with complementary traits, including those relating to plant immunity, are grown together. Cultivar mixtures have long been considered particularly effective against airborne fungal pathogens (Cowger & Mundt, 2002a; Borg *et al*., 2018) and were widely used at the end of the last century in some European countries. The use of wheat mixtures has increased in Europe in recent years, reflecting a growing interest in the beneficial effects of mixtures. In France, for instance, the percentage of the national area occupied by mixtures of bread wheat cultivars has doubled over the last four years (4.8% in 2017, 8.6% in 2018, 11.9% in 2019, and 12.2% in 2020; FranceAgriMer). The factors affecting the efficacy of cultivar mixtures against splash-dispersed diseases, such as Septoria tritici blotch (STB) on bread wheat, have also been investigated, with a combination of experimental and modelling approaches (Gigot *et al*., 2013; Vidal *et al*., 2017; Kristoffersen *et al*., 2019). The proportions of susceptible and resistant cultivars in the mixture are known to have a determinant effect on disease reduction (Finckh *et al*., 2000; Gigot *et al*., 2014; Borg *et al*., 2018). Nevertheless, most of the experimental approaches focused on the impact of mixtures on the overall canopy, with the aim of using the most resistant plants to ‘protect’ the susceptible plants, and little is known about the consequences for the most resistant cultivar. The value of cultivar mixtures for controlling diseases lies in the complementarity of the cultivar phenotypes, providing a guarantee of efficacy. However, cultivar mixtures may affect the evolution of pathogen populations, by diversifying the selection pressures exerted by their plant hosts at field scale, potentially with different effects on the different cultivars in the mixture. Little is known about pathogen evolution (changes in virulence gene frequencies) in cultivar mixtures, particularly during the early stages of resistance gene breakdown, i.e. when the frequency of virulent strains remains low.

The heterothallic ascomycete fungus *Zymoseptoria tritici*, which causes STB, is responsible for yield losses averaging 20% on susceptible bread wheat cultivars in North-Western Europe (Fones & Gurr, 2015). The development of epidemics is dependent on both splash-dispersed asexual pycnidiospores during the wheat growing period and wind-dispersed ascospores from sexual reproduction on crop residues (Figure 1a; Suffert *et al*., 2011). The potential intra-annual selective dynamics and strong genetic recombination within *Z. tritici* populations make STB a good candidate model for investigating the issues described above. Cultivars carrying qualitative and quantitative resistances have been developed and used to control STB, leading to an overall increase in the mean resistance of commercial wheat cultivars registered in France in the last decade (from 5.2 in 2015 to 6.0 in 2019, on a scale of 1 to 9, with 9 corresponding to the most resistant cultivar; ARVALIS-Institut du Végétal/CTPS). In total, 21 *Stb* genes and 89 QTLs have been identified and mapped (Brown *et al*., 2015). *Stb16q*, one of the most recent major resistance genes to have been incorporated into several cultivars by breeders, initially provided high levels of resistance against STB in both seedlings and adult plants (Ghaffary *et al*., 2012). *Stb16q* is present in 13 wheat cultivars registered in France between 2012 and 2018, including cv. Cellule (Florimond Desprez, France). Cellule was widely used in France, becoming the third most frequent cultivar grown (8% of the area under wheat in 2016), but its use then declined following a decrease in its resistance to STB. Indeed, STB symptoms were first observed on Cellule in 2015 in France (Romain Valade, ARVALIS-Institut du Végétal, pers. com.) and in 2019 in Ireland (Kildea *et al*., 2020), highlighting the beginning of a breakdown of *Stb16q* resistance.

**Figure 1.**
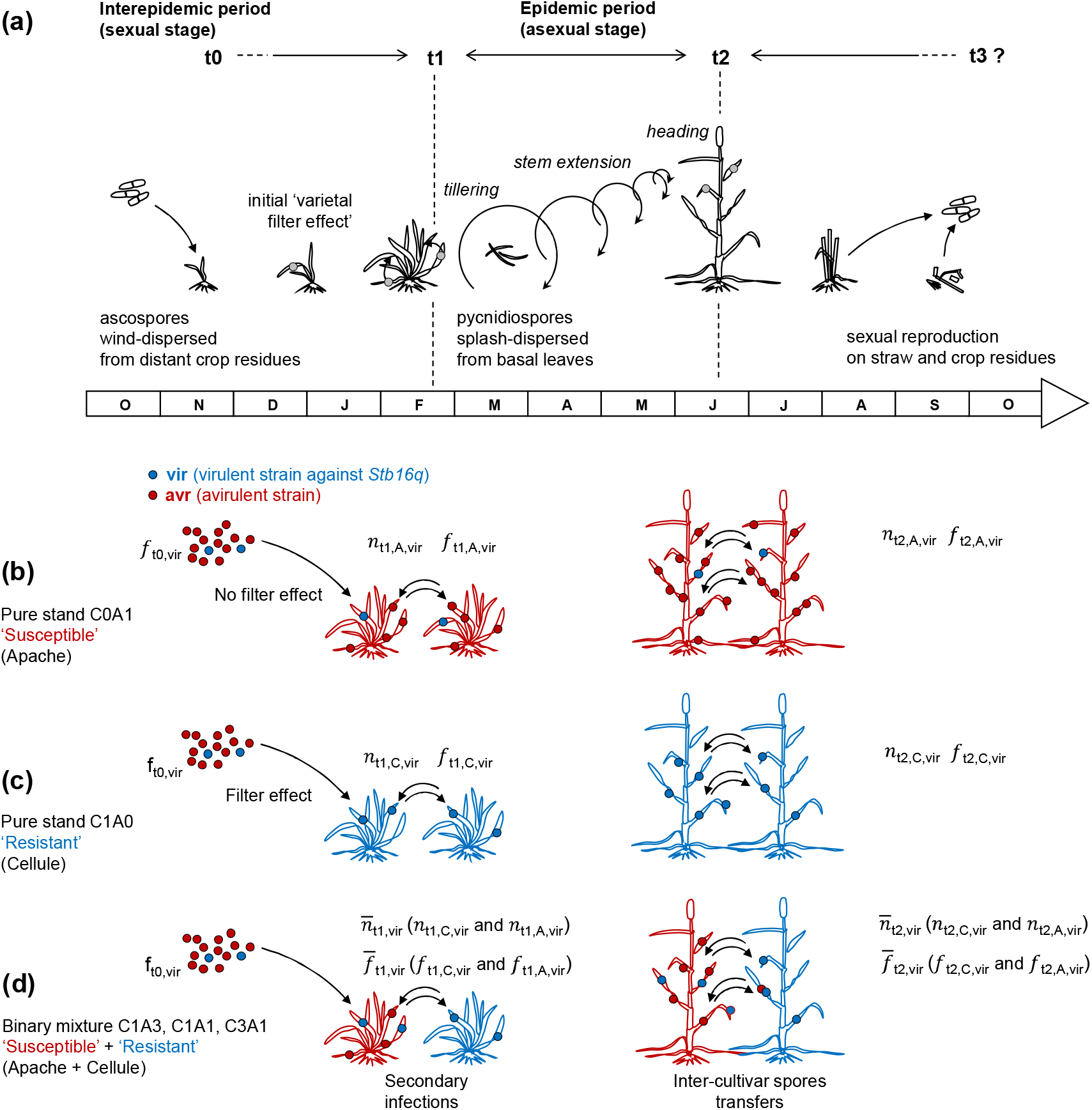
(a) Overview of the development of a Septoria tritici blotch epidemic adapted to (b, c) two pure stands and to (d) a mixture of the two wheat cultivars, one carrying the *Stb16q* resistance gene, and the other lacking this gene (cv. Cellule in proportion p_C_ and cv. Apache in proportion 1 – p_C_). The size of the pathogen population can be expressed as *n*_t,v,vir_ (size of the virulent subpopulation on the cultivar *v* ∈ [A; C] at time *t* ∈ [t1; t2]) and 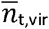 (size of the virulent subpopulation on the whole canopy at time *t*). The composition of the pathogen population is expressed as *f*_t,v,vir_ (frequency of virulent strains on cultivar *v* at *t*) and 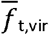 (frequency of virulent strains on the whole canopy at time *t*), with *f*_t0,vir_ the initial frequency of virulent strains in the local pathogen population.

In this epidemiological study, our goal was to investigate the capacity of mixtures to reduce disease, as a short-term epidemiological outcome, and to interfere with the resistance breakdown process, as an evolutionary outcome. In field experiments, we analysed the effect of the proportion of wheat plants carrying the qualitative resistance gene *Stb16q* within a binary wheat mixture on the size (disease severity) and composition (frequency of virulent strains) of *Z. tritici* pathogen populations during an annual epidemic.

## Materials and methods

### Overall strategy

We compared the two-year dynamics of natural *Z. tritici* populations in pure stands and in binary mixtures of wheat cultivars, from the early to late stages of a field epidemic (Figure 1a). We combined analyses of population ‘size’ based on disease severity in the field as a proxy, and of population ‘composition’ based on estimations of the proportion of virulence against *Stb16q* obtained by screening several thousands of field-collected strains in seedling assays. To this end we developed and applied an *in planta* ‘population-phenotyping’ strategy in a greenhouse. We then reinforced this strategy by ‘confirmatory individual phenotyping’ in more suitable climatic conditions in growth chamber than in greenhouse (Table S3), to increase the reliability of characterisation for some of the strains.

### Cultivars studied

This study focused on two bread wheat (*Triticum aestivum*) cultivars: Apache (‘A’; Limagrain, France) was chosen for its moderate susceptibility to STB (rated 4.5, stable over time, on a scale of 1 to 9, where 9 corresponds to the most resistant cultivar, ARVALIS-Institut du Végétal/CTPS), and Cellule (‘C’; Florimond Desprez, France), which carries the *Stb16q* resistance gene (rated 7 in 2016 vs. 5 in 2020, ARVALIS-Institut du Végétal/CTPS). Apache was the predominant cultivar in France from 2000 to 2010, accounting for 24% of the area under wheat in 2003, subsequently falling to less than 5% after 2015 (Figure 2a). Cellule, registered in 2012, became the third most frequently used French cultivar, but the area under this cultivar rapidly decreased after 2016 (Figure 2b). Both cultivars were well represented in the Parisian basin and, thus, probably played a significant role in the evolutionary trajectory of the local pathogen populations collected in 2018 and 2019.

**Figure 2.**
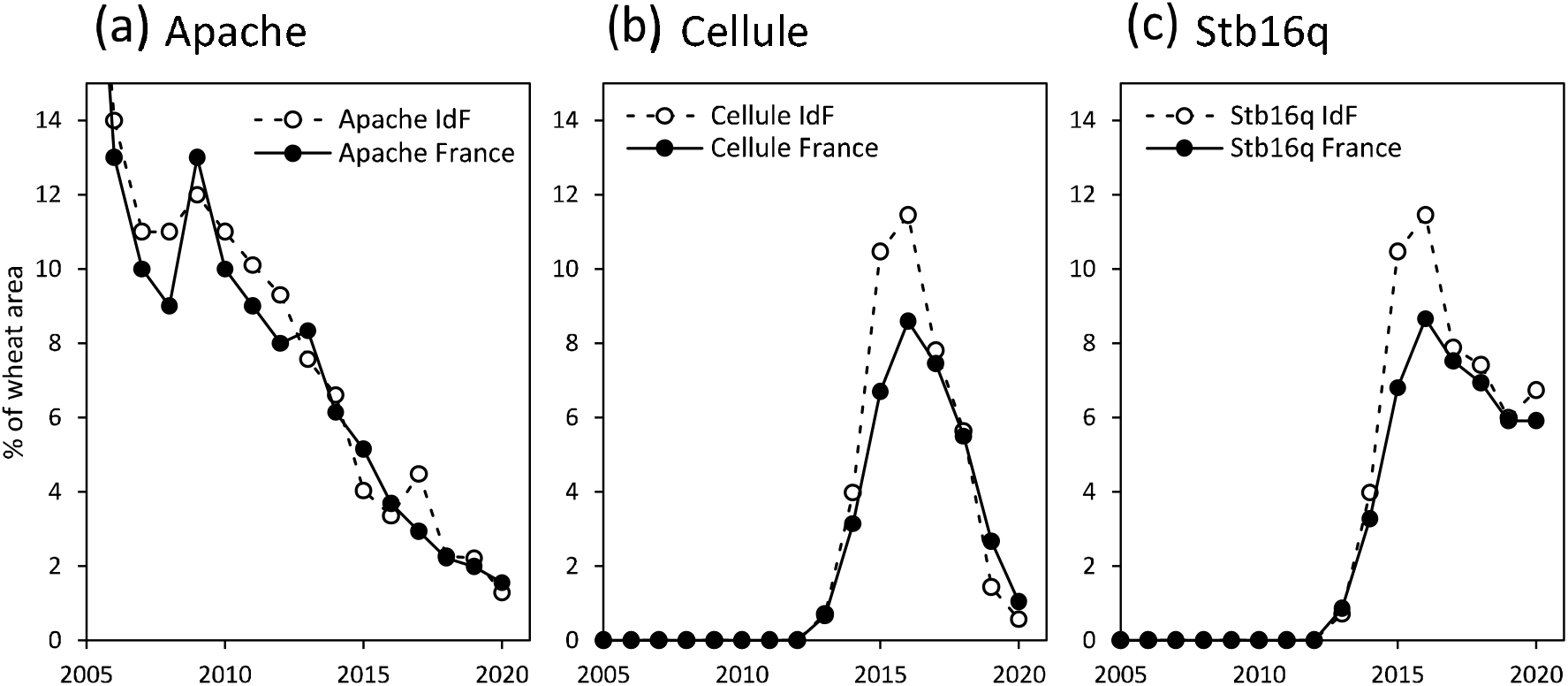
Changes in the percentage of the area under wheat sown with (a) cv. Apache, (b) cv. Cellule, and (c) the 13 cultivars carrying *Stb16q* in France (black dots) and in the Parisian basin (Région Ile-de-France ‘IdF’; white dots) from 2005 to 2020 (FranceAgriMer datasets).

### Design of the field experiment

We carried out a field experiment at the INRAE experimental station at Thiverval-Grignon (France) in the 2017-2018 and 2018-2019 seasons. A field plot was divided into three blocks (a, b and c), each consisting in five microplots (3.5 m wide x 8 m long, with an inter-row spacing of 0.175 m; Figure S1 and S2). These microplots corresponded to five randomly distributed treatments: a pure stand of wheat cv. Apache (C0A1; Figure 1b), a pure stand of wheat cv. Cellule (C1A0; Figure 1c), and three binary mixtures, with sowing proportions of Cellule (p_C_) of 0.25, 0.5 and 0.75 (C1A3, C1A1 and C3A1; Figure 1d). The wheat was sown on October 17, 2017 and November 15, 2018, at a density of 220 seeds·m^-2^. The seeds were treated with the insecticide tefluthrin at a dose of 0.2 g·kg^-1^ (Attack^®^, Syngenta France SAS) to minimize early attacks of aphids, known to transmit yellow dwarf viruses, especially BYDV. The seed lots were prepared by considering the specific weight of seeds for each cultivar (thousand-kernel-weight) to obtain the proportions of seeds required for each treatment. The sowing was performed using the different seed lots leading to a ‘randomly cultivar mixture’ at the microplot scale. Differentiation between the two cultivars — a key point of the experiment — was adapted to each growth stage: (i) at seedling stage, the identification was made according to a dye included in the seed coating (red for Cellule, green for Apache; Figure S2b); (ii) later in the season, the identification was made according to the bearded (Cellule) or unbearded (Apache) nature of the inflorescence (Figure S2e). We estimated the actual proportions of each cultivar late in the growing season, at heading, by counting spikes of each cultivar along three 1-m lines in each microplot (Figure S1, Table S1). Nitrogen fertiliser applications were adjusted by the balance-sheet method with a target yield of 7 t·ha^-1^. A single herbicide treatment was applied in the early spring. No fungicide treatment was applied.

### Characterisation of the pathogen populations size

We used disease severity (i.e., diseased leaf area) as a proxy for the size of the pathogen population (i.e., the number of asexual offspring), assuming the two traits to be correlated (Suffert *et al*., 2013). Disease severity was measured at leaf scale as the percentage of leaf area covered by pycnidia (sporulating or not). For the sake of simplicity, this measure is hereafter called “sporulating area”. Sporulating area was scored 0, 1%, 3%, 5%, 10% and then increments of 5% up to 100%. It was assessed visually early in the epidemic (time t1; Zadoks stage 15), before the expression of most of the secondary infections, and then late in the epidemic (time t2; Zadoks stage 70), just after the disease had reached the upper leaves (Figure S2). The t1 assessments of disease severity were performed on various dates (from March 5 to March 12 in 2018 and February 27 to March 18 in 2019), due to experimental constraints. We collected 10 seedlings of each cultivar from each microplot, on which disease severity was scored for the five lowest leaves. At t2, similar disease severity assessments were performed for the five upper leaves (L1 to L5 from the top to the bottom of the plant) of 10 plants of each cultivar per microplot, on May 11, 2018 and June 18, 2019. The optimal scoring window varies with epidemic dynamic. Scoring was therefore performed earlier in 2018 (intense early epidemic) than in 2019 (moderately intense late epidemic). The flag leaves (L1) were assessed a second time, in all microplots, on June 15, 2018 and June 26, 2019. The complete data set is presented in supplementary files F1-3.

A disease severity index (SEV) was calculated at the plant scale. At t1, SEV was calculated as the mean sporulating area over all non-senescent leaves. At t2, SEV was calculated as the mean sporulating area of representative leaf layers (L4 and L3 in 2018, and L3 and L2 in 2019), excluding those with a mean severity below 1% or completely senescent on more than two microplots.

A mean disease severity index (SEV_v_) was then calculated for each cultivar *v* for each treatment. An overall mean disease severity index 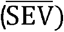 was also calculated at the whole-canopy scale for each treatment, using SEV_v_ weighted by the proportions of the cultivars (p_C_ for Cellule and 1 – p_C_ for Apache), as follows:

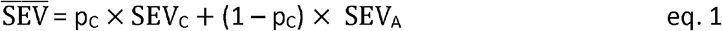

We assessed the effect of treatment on the disease severity index (SEV) at t1 and t2, and analysed changes in this effect over the course of each epidemic. To this end, SEV was transformed by ordered quantile normalisation (*bestNormalize* R package). Its variance was partitioned into sources attributable to the following factors and their interactions, in pairs: cultivars (A; C), treatments (C0A1; C1A3; C1A1; C3A1; C1A0), blocks (a; b; c), epidemic stages (t1; t2) and years (2018; 2019), according to the generalised linear model. The model was adjusted by removing non-significant interactions one-by-one. For SEV_v_ and 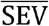, standard errors were corrected by taking into account the error propagation of formulas.

The effect of treatment on SEV_A_ and SEV_C_ at each stage of each year was first assessed with a Tukey HSD post-hoc test for multiple pairwise comparisons on estimated marginal means (*emmeans* and *multcomp* packages of R). The variance of the overall disease severity index 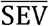 was then estimated by calculating the standard error. The effect of cultivar mixtures on 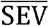 was evaluated by plotting the ‘expected’ 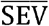 (assuming an absence of mixture effect) as a linear function of the severity in each pure stand weighted by Cellule proportions. All data analyses were performed with R software (v4.0.2 R Core Team 2012).

### Characterisation of the pathogen populations composition

#### Sampling of strains in the field

Strains of *Z. tritici* were collected at t1 and t2 in 2018 and 2019, to constitute groups of 30 strains per cultivar and per microplot (total of 3060 strains; Table 1 and supplementary file F4). The sampling effort was doubled in the pure stands of the susceptible cultivar at t1 in 2018 and 2019 (2 × 180 strains). The population present on cv. Apache early in the epidemic (t1), which may be considered a representative sample of the primary inoculum population, was used to estimate the initial (t0) frequency of virulent strains (*f*_t0_,_vir_; Figure 1b).

**Table 1.** Number of *Z. tritici* strains sampled on each cultivar (‘A’ for Apache, ‘C’ for Cellule) for each treatment (C0A1, C1A3, C1A1, C3A1 and C1A0), at the early (t1) and late (t2) stages of the 2018 and 2019 epidemics, and phenotyped to determine virulence status.

| Year | Stage | COA1 | C1A3 |  | C1A1 |  | C3A1 |  | C1A0 | Total |
| --- | --- | --- | --- | --- | --- | --- | --- | --- | --- | --- |
|  |  | A | C | A | C | A | C | A | C |  |
| 2018 | t1<sup>1</sup> | 179 | 90 | 90 | 90 | 90 | 90 | 87 | 90 | 806 |
|  | t2<sup>2</sup> | 90 | 90 | 90 | 89 | 90 | 90 | 90 | 90 | 719 |
| 2019 | t1<sup>3</sup> | 180 | 82 | 88 | 88 | 84 | 90 | 89 | 90 | 791 |
|  | t2<sup>4</sup> | 90 | 90 | 90 | 90 | 90 | 90 | 89 | 90 | 719 |

<sup>1</sup> collected from March 5<sup>th</sup> to 12<sup>th</sup>; <sup>2</sup> June 13<sup>th</sup>; <sup>3</sup> from February 27<sup>th</sup> to March 13<sup>th</sup>; <sup>4</sup> June 25<sup>th</sup>

We ensured that 30 strains were obtained per cultivar from each microplot, by sampling the 40 lowest leaves of seedlings at t1 and the flag leaves of adult plants at t2. These samples were placed in a humid chamber to promote the extrusion of cirrhi. One cirrhus per leaf was picked with a pin and streaked onto a Petri dish containing PDA (potato dextrose agar, 39 g·L^-1^), which was then incubated at 18°C for more than five days in the dark. A few blastospores from a single colony were then picked and spread on a new plate. This operation was repeated twice to obtain pure single-spore strains. Each strain was finally scraped off the plate, to obtain a substantial mass of blastopores (150 mg), which was deposited at the bottom of cryotubes and stored at -80°C. At t1 in 2018, we sometimes had to collect more than one strain per leaf to counterbalance the low level of severity on certain microplots or the degradation/senescence of the diseased leaves. In both situations, we picked cirrhi from two to three distinct, distant lesions present on the same leaf.

#### Phenotyping strategy

We characterised the *Z. tritici* strains for ‘virulence’ (defined by Niks *et al*. (2011) as “the capacity of a pathogen to infect a plant with one or more major genes for resistance”) against *Stb16q* and their ‘aggressiveness’ (the quantitative term of pathogenicity on a susceptible host; Lannou, 2012). We developed a ‘population-phenotyping’ method for use on cv. Cellule and cv. Apache seedlings in the greenhouse (Figure S3), which allowed us to characterise individually a large number of strains while accepting some imprecisions or individual errors quantified *a posteriori*. This approach was complemented by ‘confirmatory individual phenotyping’ in growth chambers, in which we characterised several dozens of strains that were collected from sporulating lesions on Cellule in the field but appeared avirulent during population-phenotyping. This made it possible to determine the rate of type II error (false avirulence) for population-phenotyping.

#### Population-phenotyping

Population-phenotyping was performed from April 2018 to August 2020, by splitting the 3060 strains into 32 batches of 102 (t1) or 93 (t2) strains, which were simultaneously tested in the greenhouse (Figure 3). Each batch was balanced to contain similar numbers of strains from each cultivar, treatment, and block. Eight virulent strains were added to each batch as control strains. Wheat seeds (cv. Apache or Cellule) were sown in 0.4-litre pots, to obtain three 18-day-old seedlings per pot. The pots were kept in the greenhouse under natural daylight supplemented with 400 W sodium vapour lamps to homogenise the lighting conditions. The temperature was kept below 20°C during the 15-h light period and above 15°C during the 9-h dark period. Relative humidity was kept at a mean value of 68 ± 9%. The thermal time at which disease severity was assessed was calculated for each batch (Figure S5).

**Figure 3.**
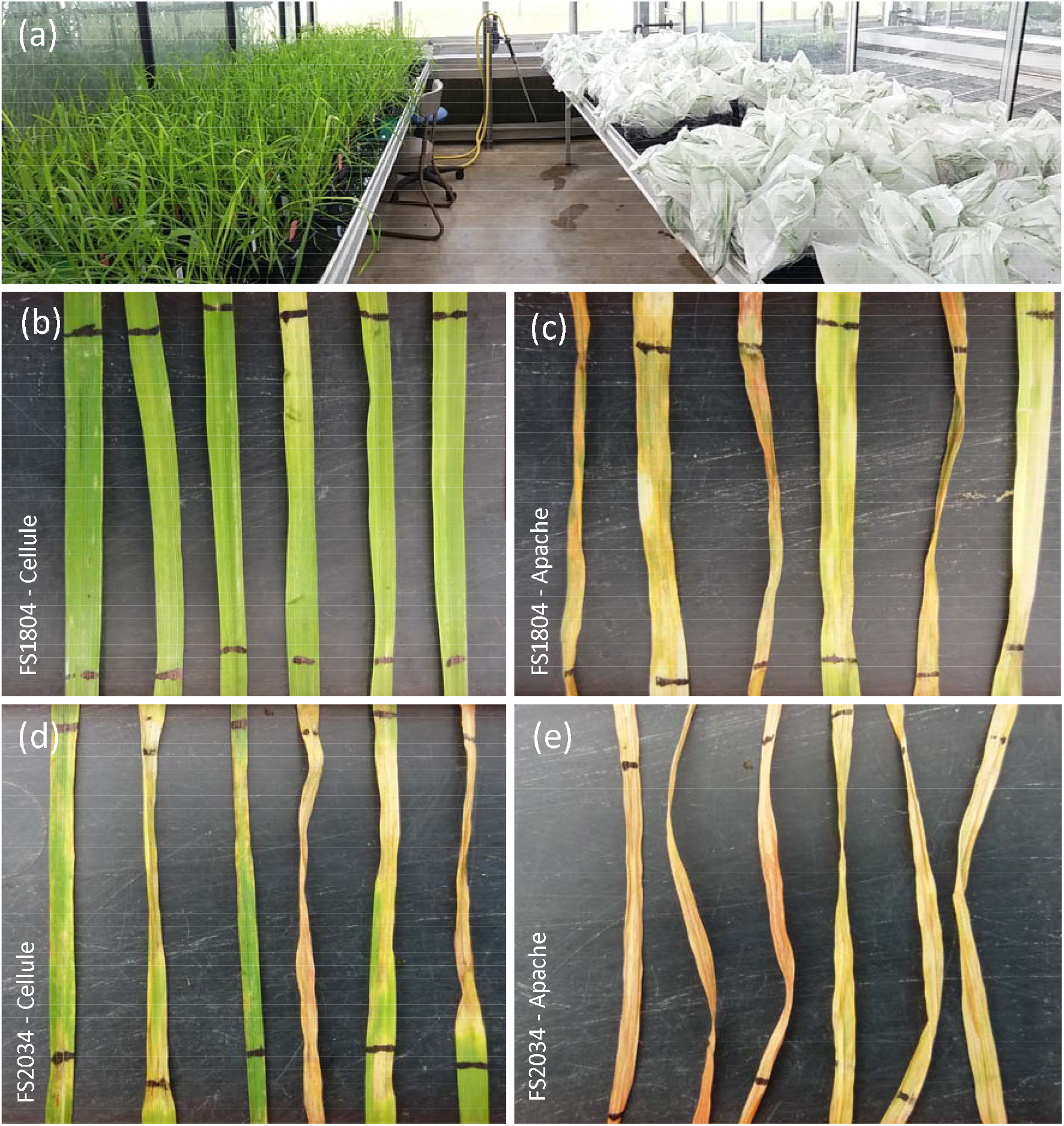
(a) Example of two 102-strain batches, one obtained just after inoculation (right) and the other one week later (left). (b-e) Typical incompatible-compatible interaction between the FS1804 (avirulent against *Stb16q*) and FS2034 (virulent against *Stb16q*) strains on six Cellule and Apache leaves (three second leaves on the left and three third leaves on the right).

We simplified the classical method based on accurate adjustment of a blastospore suspension, by considering that the differences in disease severity induced by the differences in concentration within the range 10^5^ to 5.0 10^5^ spores·mL^-1^ (see Figure 8 in Suffert *et al*., 2018) were acceptable for our objective. We thus used non-adjusted suspensions in this range of concentrations. For each strain, we diluted the blastospores recovered directly with a 10 μL inoculating loop (about 10 mg in the half-filled central ring; Sarstedt, Nümbrecht, Germany) from the freshly thawed content of the cryotube preserved at -80°C, in 30 mL of water. Two series of independent tests showed that this process provided a good compromise between robustness and repeatability (Figure S4). Two drops of surfactant (0.1% v/v of Tween 20; Sigma, France) were added to each suspension, to increase its adhesion to leaves. All suspensions were prepared the day before inoculation and stored at 5°C. A cotton bud was used to inoculate a 5 cm mid-length leaf section by two back-and-forth passages over the second and third leaves from the bottom on seedlings previously described. This way, each strain was tested individually on two leaves of the three seedlings of the pot (6 replicates), for each cultivar. Each pot was then enclosed in sealed transparent polyethylene bags moistened with distilled water for 72 h, to promote infection.

Disease severity was assessed visually 21 days post inoculation (mean thermal time 434.8°C·days for the 32 batches; Figure S5), as the percentage of the inoculated leaf surface covered by pycnidia (sporulating area assessed, from 0 to 100%, in increments of 5%). The complete data set is presented in supplementary file F5. A strain was considered virulent (‘vir’) against *Stb16q* when pycnidia were observed on at least one of the six leaves of Cellule (at least 5% of sporulating area on the leaf, the minimum for visual quantification in our experimental conditions). The strains were otherwise considered avirulent (‘avr’). The frequency of virulent strains *f*_t,v,vir_ (t ∈ [t1; t2], v ∈ [A; C]) was calculated for each cultivar in each treatment (Figure 1b-d). The effect of the treatment on the number of virulent individuals was assessed with a 𝒳^2^ test, or with Fisher’s exact test when the expected numbers were small, with Bonferroni correction for pairwise comparison. We analysed the changes in *f*_t,v,vir_ from t1 to t2, assessing the year effect with a test of equal or given proportions (*stats* package of R). The initial frequency of virulent strains in the local pathogen population (*f*_t0,vir_; Figure 1b-d) in 2018 and 2019 was estimated by characterising the 180 strains from the pure stand of cv. Apache. We postulated that in this treatment the initial ‘filter effect’ exerted on a strain according to its virulence status with respect to *Stb16q* was negligible (*f*_t1,A,vir_ = *f*_t0,vir_ in C0A1; Figure 1b).

We calculated the aggressiveness (AG) and conditional aggressiveness (AGc), defined as the amount of disease “conditional on the individuals being infected” (McRoberts *et al*., 2003). AG was estimated as the mean sporulating area on all leaves of the susceptible cultivar, whereas AGc was calculated on symptomatic leaves only, to take into account the absence of symptoms on some leaves (due to the less suitable climatic conditions in greenhouse than in growth chamber; Table S3). For a given strain, AGc was thus calculated under the condition that at least three of the six inoculated leaves presented pycnidia. Furthermore, we assessed the possible fitness cost of virulence against *Stb16q*, by evaluating the effect of virulence on aggressiveness. Because of the non-normality of data and the shape of the distribution curve, we used a generalised linear model adapted to a Poisson distribution, followed by a Tukey HSD post-hoc comparison (*car R package*). The variance of each aggressiveness trait (AG and AGc) was partitioned into sources attributable to the following factors and their interactions, in pairs: the virulence status of the strain (vir; avir), treatment (C0A1; C1A3; C1A1; C3A1; C1A0), the cultivar of origin (A; C), year (2018;2019), epidemic stage (t1; t2), block (a; b; c), and batch (1-32).

#### Confirmatory individual phenotyping

A second round of phenotyping was applied to 47 strains that did not fulfil the requirement to be considered virulent against *Stb16q* during the population-phenotyping in greenhouse, despite having been isolated from sporulating lesions on Cellule in 2018 (see the Results section). For this purpose, we used Cellule (C) and two isogenic lines of the wheat cultivar Chinese Spring without *Stb6* (naturally present in Chinese Spring, Chartrain *et al*., 2005), one of which carried *Stb16q* (CS-*Stb16q*) whereas the other did not (CS). The strains were split into three batches and tested individually in a growth chamber (Figure S3). A virulent and an avirulent strain were added to each batch as controls. Inoculum suspensions were adjusted to a density of 5.0 10^5^ spores·mL^-1^ with a Malassez cell. Leaves were inoculated by applying the suspension with a paintbrush, on the second leaf from the bottom of nine seedlings of each cultivar. The temperature in the growth chamber was kept below 20°C during the 15-h light period and above 15°C during the 9-h dark period. Relative humidity was maintained at a mean of 88.5 ± 1.2%. Disease severity was assessed visually 14, 20 and 26 dpi (day post inoculation), as the percentage sporulating area. The virulent, avirulent or non-pathogenic nature of each strain was determined by calculating the mean aggressiveness (mean percentage of sporulating area on a leaf obtained across the nine replicates) and compared it to a classical 10% of sporulating area threshold (Thierry Marcel, INRAE BIOGER, pers. com.) rather than 5% in greenhouse (less suitable climatic conditions; Table S3). A strain was considered (i) virulent against *Stb16q* if the mean aggressiveness was equal or higher than this threshold on both CS-*Stb16q* and Cellule, (ii) avirulent if it was equal or higher than this threshold on CS only, or (iii) non-pathogenic if it was below this threshold on the three cultivars (Figure S10). The mean aggressiveness of strains was compared between CS-*Stb16q* and Cellule with a Wilcoxon test. The effect of treatment on the number of avirulent strains was assessed with Fisher’s exact test.

### Overall impact of cultivar mixtures on the dynamics of virulent subpopulations

We calculated the percentage of the diseased leaf area occupied by virulent strains at t2 (*n*_t2,v,vir_; v ∈ [A; C]; Figure 1b-d), as a proxy for the size of the virulent subpopulations responsible for the last asexual disease cycle. This dimensionless variable was obtained by multiplying the disease severity index (SEV_v_) on the flag leaves of each cultivar in each microplot by the frequency of virulent strains estimated at t2 (*f*_t2,v,vir_). The mean size of the virulent subpopulations 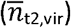 and the mean frequency of virulent strains 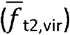 were calculated at the whole-canopy scale for each treatment (Figure 1b-d), as for overall disease severity index (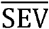, eq. 1). The effect of treatment on *n*_t2,v,vir_ and 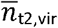 was assessed in a Kruskal-Wallis test.

## Results

### Impact of cultivar mixtures on the dynamics of the epidemic and the overall size of pathogen populations

Disease was generally less severe in the mixtures than in the two pure stands late in the epidemic, when the whole canopy was considered, and on both cultivars relative to their pure stands. This can be interpreted as a reciprocal ‘protective’ effect.

At t1 in 2018, no significant difference in mean disease severity index (SEV_v_) was observed between pure stands and mixtures, regardless of the cultivar considered, except for Apache in C1A1 (*p* = 0.987, *p* = 0.042 and *p* = 0.155 for Apache and *p* = 0.172, *p* = 0.388, and *p* = 0.065 for Cellule in C1A3, C1A1, and C3A1, respectively; Figure 4). In mixtures, Apache was in average 1.6 times more attacked than Cellule, consistent with their resistance ratings. The modelling results showed that the variance of disease severity index (SEV) was not attributable to the block factor (*p* = 0.090). Overall disease severity index 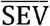 decreased with the proportion of Cellule. At t1 in 2019, SEV_A_ and SEV_C_ were lower than in 2018. In mixtures, Apache was in average 2.8 times more attacked than Cellule. In pure stands, SEV_A_ scored 17% and SEV_c_ scored 4% (data none shown). 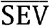 decreased with the proportion of Cellule.

**Figure 4.**
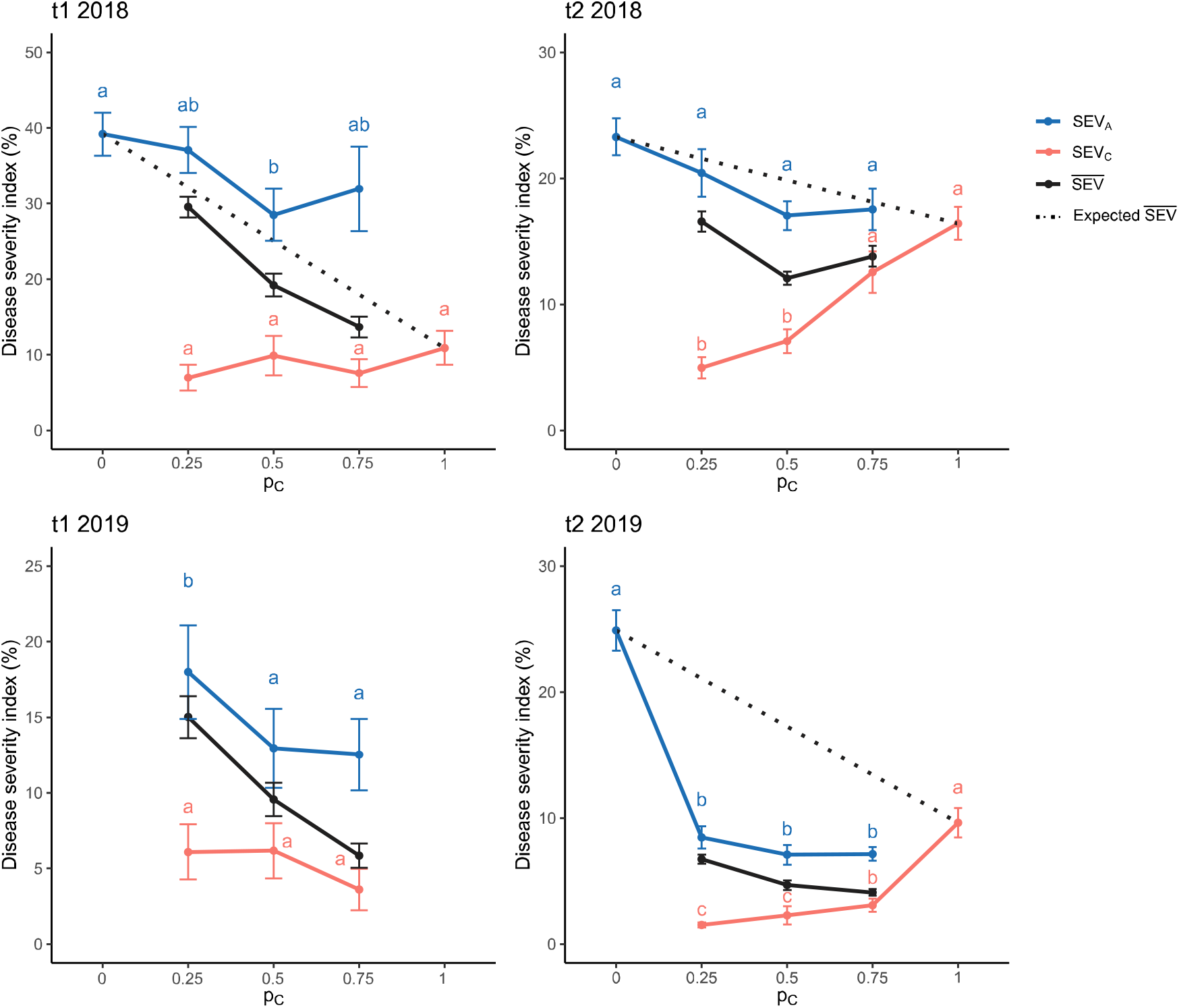
Mean disease severity index SEV_v_ (mean percentage of the leaf area covered by pycnidia on cultivar *v*) according to the proportion of cv. Cellule (p_C_) in the canopy. SEV_A_ (green solid line) and SEV_C_ (red solid line) were calculated from disease assessments in the field. Bars represent the standard error. Significant differences in SEV_A_ and SEV_C_ between p_c_ are indicated by letters (generalised linear model followed by a Tuckey HSD post-hoc test. The overall severity index 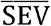 (black solid line) was calculated at the whole-canopy scale as the mean of SEV_A_ and SEV_C_ weighted by the proportion of each cultivar. The ‘expected’ 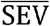 (in the absence of a mixture effect) was plotted as a linear function of the severity in each pure stand weighted by p_C_ (black dotted lines). SEV_A_, SEV_C_ and the expected 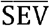 in pure stands at t1 in 2019 are not shown because disease severity was assessed one week earlier than in mixtures.

At t2 in 2018, SEV_v_ differed significantly between pure stands and mixtures for Cellule, but not for Apache (*p* = 0.397, *p* = 0.148, and *p* = 0.357 for Apache and *p* < 0.001, *p* < 0.001, and *p* = 0.089 for Cellule in C1A3, C1A1, and C3A1, respectively; Figure 4). In pure stands, Apache was in average 1.5 times more attacked than Cellule. In mixtures, SEV_A_ was in average reduced by 20% and SEV_C_, to a greater extent, reduced by 48% compared to the respective pure stands. At the whole-canopy scale, 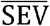 was lower in the three mixtures than in pure stands. At t2 in 2019, SEV_v_ differed significantly between pure stands and mixtures for both cultivars, this difference being more significant than in 2018 (*p* < 0.0001 for Apache and *p* < 0.0001 for Cellule in all C1A3, C1A1, and C3A1). In pure stands, Apache was 2.5 times more attacked than Cellule. In mixtures, SEV_A_ was in average reduced by 69% and SEV_C_ reduced by 73% compared to the respective pure stands. The decreases in SEV_A_, SEV_C_, and 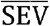 in mixtures relative to pure stands was more pronounced in 2019 than in 2018.

The overall effect of treatment was illustrated by plotting the expected 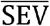 as a linear function of the severity in each pure stand weighted by Cellule proportions (p_C_; Figure 4), which accounts only for ‘dilution’ in mixtures. The actual and expected 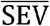 curves were close at t1 in 2018 whereas the actual 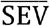 was below the expected 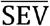 at t2 for the two years considered. This highlights the overall absence of a mixture effect on disease dynamics at t1, at least in 2018, and the strong mixture effect at t2 for both years. The efficacy of the various treatments relative to pure stands was 19%, 40%, 22% in 2018 and 67%, 71%, 69% in 2019, for C1A3, C1A1, and C3A1, respectively.

### Impact of mixtures on the composition of pathogen populations

In total, 3035 of the 3060 *Z. tritici* strains sampled during the two years were characterised for virulence and aggressiveness, by inoculating 36420 leaves of wheat seedlings in the greenhouse. The cultivar mixtures increased the frequency of virulent strains from the early (t1) to late (t2) stages in the two epidemic years.

The frequency of virulence at t1 (*f*_t1,A,vir_) was estimated on Apache in pure stands at 0.13 in 2018 and 0.14 in 2019 (Figure 5). These frequencies are not significantly different (χ2 test, *p* = 0.791), and the two years may therefore be considered as ‘replicates’. The frequency of virulent strains on Apache did not differ significantly between treatments at t1, confirming the absence of a mixture effect early in the epidemic. The frequency of virulent strains on Apache at t2 (*f*_t2,A,vir_) in pure stands was 0.08 in 2018 and 0.18 in 2019, and was not significantly lower than that at t1 (test of equal or given proportions, *p* = 0.244). Importantly, in mixtures, *f*_t2,A,vir_ increased significantly with p_C_ both years.

**Figure 5.**
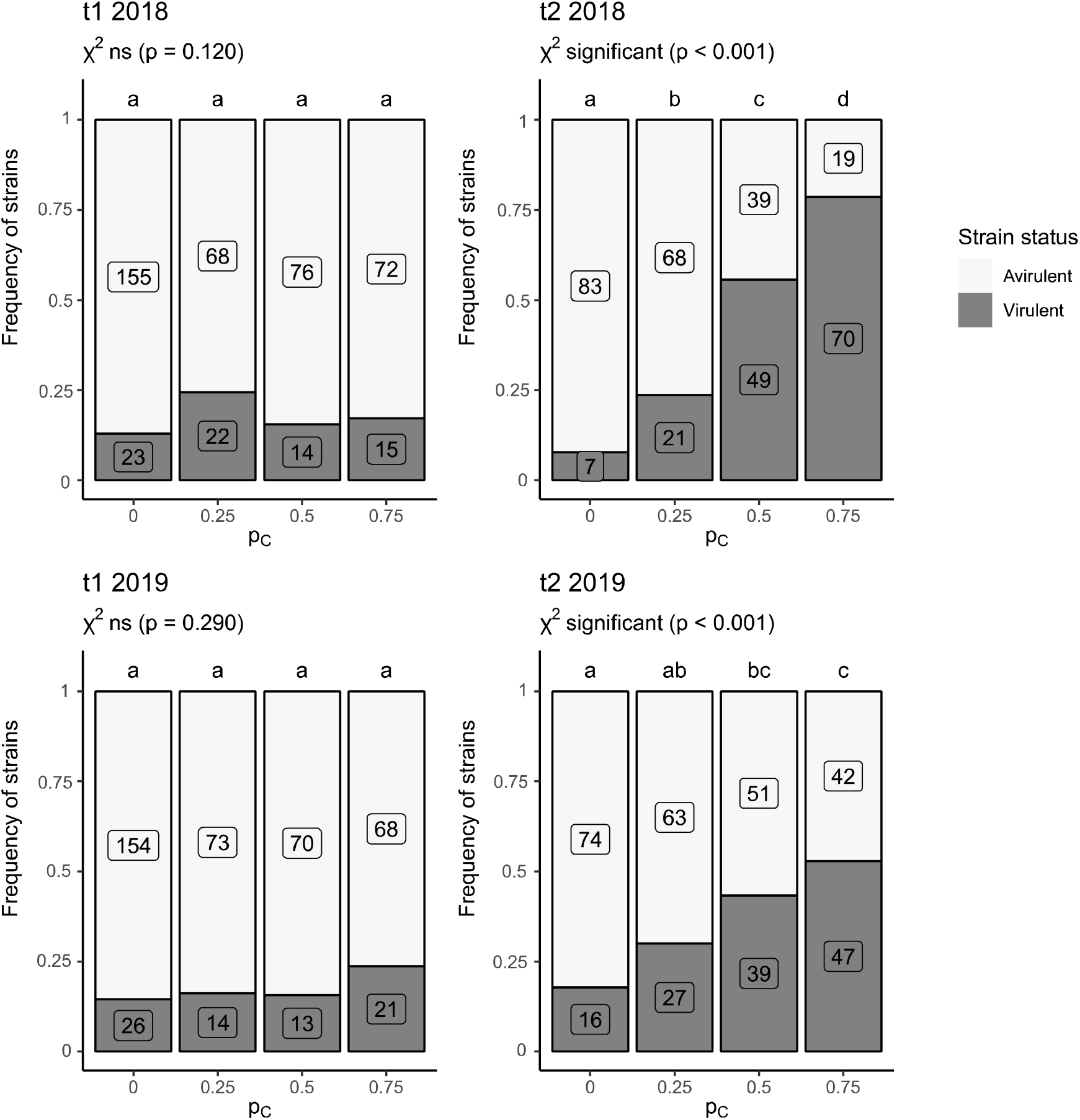
Frequency of *Z. tritici* strains considered virulent (grey bars) against *Stb16q* (i.e., causing sporulating area on cv. Cellule) and avirulent (white bars) among the populations collected on cv. Apache in pure stands (p_C_ = 0) and mixtures (p_C_ = 0.25, 0.5 and 0.75 for C1A3, C1A1 and C3A1, respectively) at the early (t1) and late (t2) stages of the 2018 and 2019 epidemics. The number of virulent and avirulent strains are indicated in grey and white boxes, respectively. The effect of treatment (pure stands and cultivar mixtures; letters at the top of each bar) on the number of virulent strains was assessed by performing a χ² test with Bonferroni correction for pairwise comparison.

Very few strains were classified as non-pathogenic: 0.3% (5/1525) in 2018 and 0.1% (2/1510) in 2019 (Figure S6 and S7). Most of the strains (>95%) induced pycnidia on Apache, with no significant difference between treatments. The percentage of strains that did not induce pycnidia on Apache was low: 4% (31/719) and 3% (23/710) for those collected on Cellule in 2018 and 2019, respectively, and 6% (50/806) and 5% (38/800) for those collected on Apache in 2018 and in 2019, respectively. Strains collected on Cellule were expected to be virulent, as a consequence of the ‘filter effect’ exerted by *Stb16q* (Figure 1c). However, 7% of these strains (52/719) in 2018 and 10% (74/710) in 2019 (Figure 6) did not induce pycnidia on Cellule during phenotyping. The proportion of these strains, considered avirulent after population-phenotyping, did not differ significantly between treatments. Only 2% of the strains (27/1525 in 2018 and 20/1510 in 2019, distributed across all treatments) induced pycnidia on Cellule but not on Apache.

**Figure 6.**
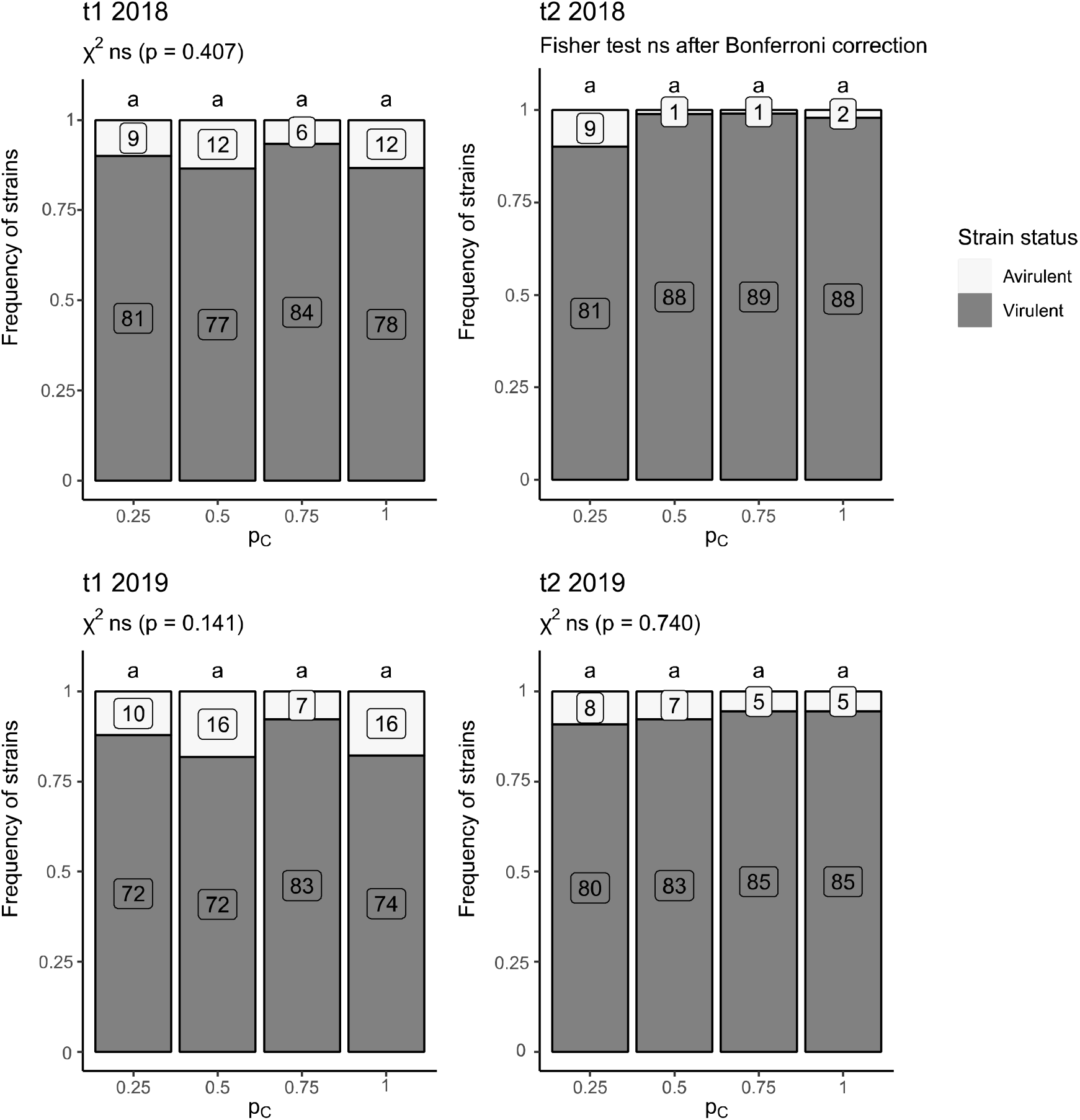
Frequency of *Z. tritici* strains considered virulent (grey bars) against *Stb16q* (i.e., causing sporulating area on cv. Cellule) and avirulent (white bars) among the populations collected on cv. Cellule in pure stands (p_C_ = 1) and mixtures (p_C_ = 0.25, 0.5 and 0.75 for C1A3, C1A1 and C3A1, respectively) at the early (t1) and late (t2) stages of the 2018 and 2019 epidemics. The number of virulent and avirulent strains are indicated in grey and white boxes, respectively. The effect of treatment (pure stands and cultivar mixtures; letters at the top of each bar) on the number of virulent strains was assessed by performing a χ^2^ test, or a Fisher’s exact test when the expected numbers were small, with Bonferroni correction for pairwise comparison.

Confirmatory individual phenotyping showed that 51% (24/47) of the strains tested were truly avirulent, 34% (16/47) were virulent, and the results were dubious for 15% (7/47). The overall percentage of ‘false avirulent’ strains collected on Cellule in 2018 was thus estimated at a maximum of 3% (23/719). All the truly avirulent strains were collected at t1 (Table S2). Five of the strains yielding dubious results that induced pycnidia on CS-*Stb16q* were unable to infect Cellule, probably due to another source of resistance not present in CS. Similarly, two strains that induced pycnidia on Cellule were unable to infect CS-*Stb16q*. All 47 strains induced pycnidia on CS, confirming their pathogenicity.

Virulent strains were significantly more aggressive on CS-*Stb16q* than on Cellule (Wilcoxon test, *p* < 0.0001; Figure S9). No significant difference in aggressiveness (neither in AG nor AGc; *p* = 0.054 and *p* = 0.597, respectively; Figure S8) was detected between the avirulent and virulent strains that were pathogenic on Apache (2886/3035).

### Impact of mixtures on the dynamics of the virulent subpopulation

At whole-canopy scale, cultivar proportions modulated the decreases in the size of the virulent subpopulation and in the frequency of virulent strains relative to the pure stand of the cultivar carrying *Stb16q* (Cellule).

In 2018, the mean size of the virulent subpopulation 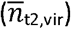 was significantly higher in pure stands of Cellule than in pure stands on Apache. However, 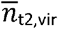 was not significantly different in mixtures than in pure stands of Cellule or Apache (Figure 7a). 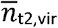 tended to increased with the proportion of Cellule (p_C_). In 2019, the three mixtures significantly decreased 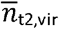 relative to the pure stand of Cellule (Figure 7b), whereas the difference was not significant relative to the pure stand of Apache. The figure 7b shows a correlation of means and variances, assuming that mixture may both reduce and stabilize the virulent subpopulation size compare to pure stands. In 2019, 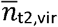 in mixtures was lower than in pure stand of Cellule by a factor of about 5 and lower than that in the pure stand of Apache by a factor of 2. This difference can be explained by the very low frequency of virulent strains on Apache (*f*_t2,A,vir_) being compensated in 2018, and, to a greater extent in 2019, by the size of the pathogen population on Apache (SEV_A_), resulting in a larger number of virulent strains 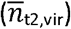. A dissociated analysis of each varietal component of the mixture can provide insight on this point. On Apache, the size of the virulent subpopulation (*n*_t2,A,vir_) was not significantly different between pure stands and mixtures in 2018 (*p* = 0.200, Kruskal-Wallis test; Figure S11a) or 2019 (*p* = 0.578, Kruskal-Wallis test; Figure S11c). On Cellule, the size of the virulent subpopulation (*n*_t2,C,vir_) increased significantly with p_C_ in 2018 (*p* = 0.033) and 2019 (*p* = 0.024). Thus, only *n*_t2,C,vir_ contributes to the differences at whole-canopy scale.

**Figure 7.**
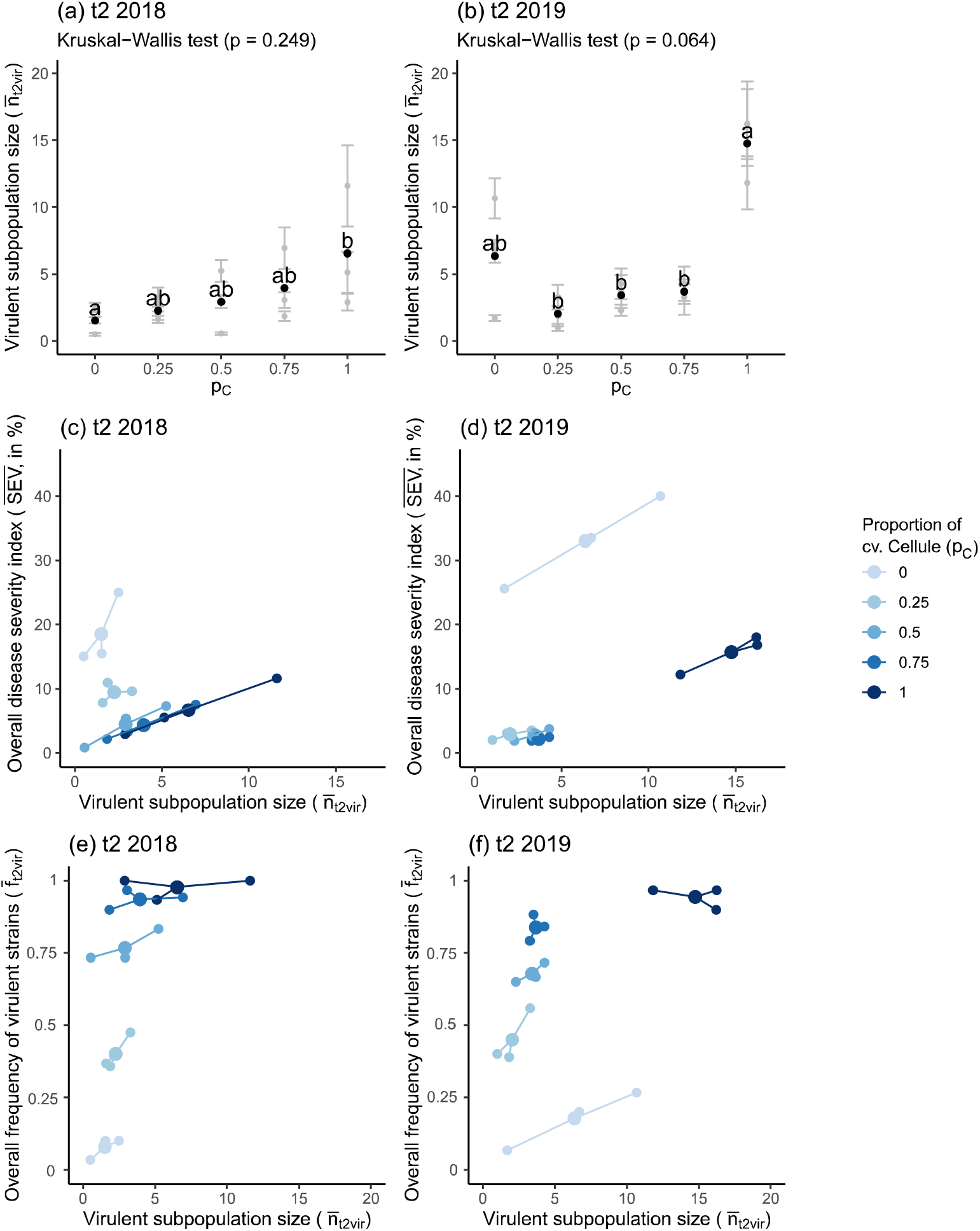
Impact of mixtures on different indicators of the *Z. tritici* virulent subpopulations and their relationships at whole-canopy scale. (a, b) Size of the virulent subpopulations at t2 (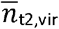 dimensionless variable; black dots) according to the proportion of cv. Cellule (p_C_) in 2018 and 2019. Grey dots indicate the mean value for each block and black dots the mean for each treatment. Bars represent the standard error within each block. Error propagation was taken into account. (c, d) Disease severity assessed at the whole-canopy scale at t2 (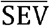, determined as the percentage of the leaf area covered with pycnidia on flag leaves) according to the size of the virulent subpopulation 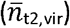. (e, f) Frequency of virulent strains assessed at whole-canopy scale at t2 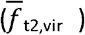 according to the size of the virulent subpopulations in 2018 and in 2019. Small dots indicate the mean value for each block, and large dots the mean value for each treatment. Different shades of blue represent the proportion of cv. Cellule (p_C_).

We examined the relationship between the indicators investigated. We first focused on the relationship between the overall disease severity 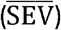— an indicator of the efficacy of the mixtures — and the overall size of the virulent subpopulation 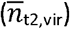, an indicator of *Stb16q* breakdown dynamics and, thus, a proxy for the impact of the mixtures on resistance durability (Figure 7c and 7d). In both years, the overall trend was similar for the virulent subpopulation: 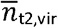 increased with the proportion of Cellule (p_C_) whereas 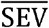 decreased. In 2018, two mixtures (C1A1 and C3A1) displayed decreases for both 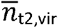 and 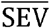 relative to the pure stands of Cellule and Apache. In the C1A3 mixture, 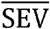 was slightly higher with a lower level of 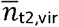 than in other mixtures. The impact of mixtures was greater in 2019, with all mixtures decreasing both 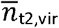 and 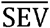, which was, on average, lower than in pure stands of Apache by a factor of 10 and lower than that in pure stands of Cellule by a factor of 5. We then focused on the relationship between the mean frequency of virulent strains 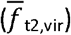 and the overall size of the virulent subpopulation 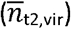 (Figure 7e and 7f). Both years, 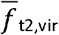 and 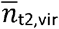 increased with p_C_, whose quantitative impact was generally high for 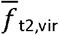 but relatively limited for 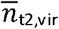, especially in 2018 for mixtures (Figure 7e and 7f).

Finally, regardless of the indicator considered 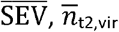 or 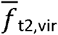 and of relationship trends, the optimal p_C_ appeared to be about 0.5. However, our results suggest that moving away from this optimal proportion would have different effects on the three indicators.

## Discussion

We found that pathogen population size, estimated on each cultivar but also at the whole-canopy scale, was smaller for mixtures than for pure stands at the end of epidemics. In addition we established that mixtures reduced the size of the virulent subpopulation and increased the frequency of strains displaying virulence against *Stb16q*, relative to pure stands of Cellule (thereafter referred as the ‘resistant’ cultivar).

### A method suitable for studies at population scale

Molecular tools are not always available for monitoring the emergence of virulence. This complicates the quantification of breakdown dynamics for recently deployed resistance genes. Our *in planta* ‘population-phenotyping’, useful for this purpose, was twice as efficient as the classical method (Figure S3). The various experimental adjustments we made to inoculum preparation, climatic conditions in the greenhouse, visual assessments and the small number of biological replicates (Table S3) introduced some uncertainty, potentially leading to diagnostic errors concerning virulence status at an individual scale (‘false avirulence’ rate estimated at 3%). However, this uncertainty had only a small impact at population scale.

### *Pathogenicity on Cellule vs. virulence against* Stb16q

We assumed that a strain pathogenic on cultivar Cellule was virulent against *Stb16q* and that strains non-pathogenic on this cultivar were avirulent. However, as the genetic background of the Cellule cultivar probably includes other sources of resistance, an absence of symptoms on Cellule is not sufficient to demonstrate avirulence against *Stb16q*. The confirmatory individual phenotyping data provide some perspective concerning this approximation: five of the seven strains inducing pycnidia on CS*-Stb16q* were unable to infect Cellule, and two of the strains pathogenic on Cellule were unable to develop sufficient pycnidia on CS-*Stb16q* to exceed the 10% threshold for virulence. The rarity of such situations (<1%) suggests that ability to induce pycnidia on Cellule is a reasonably indicator of virulence against *Stb16q*. Finally, we estimated that the virulence status established based on population-phenotyping of the Cellule response was correct in 97% of cases.

### The cultivars displayed reciprocal protection

Early in the epidemic, most of the time, the size of the pathogen population in the whole canopy 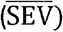 and on both susceptible (SEV_A_) and resistant (SEV_c_) cultivars was not significantly different between mixtures and the corresponding pure stands (Figure 4). This is an expected result, as mixture effects were reported to increase during asexual cycles (Cowger & Mundt, 2002a; Gigot *et al*., 2013). Late in the epidemic, cultivar mixtures reduced 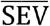 confirming the potential of wheat mixtures to limit the dynamics of STB epidemics (Kristoffersen *et al*., 2019). In our experiment, mixtures reduced 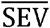 by a mean of 27% in 2018 and 69% in 2019. The size of the pathogen population was lower in mixtures than in the corresponding pure stands only on the resistant cultivar in 2018 and on both cultivars in 2019. The relative differences in SEV_v_ between mixtures and pure stands ranged from 13% to 26% in 2018 and from 64% to 72% in 2019 for the susceptible cultivar, and from 19% to 69% in 2018 and from 70% to 80% in 2019 for the resistant cultivar. Gigot *et al*. (2013) and Vidal *et al*. (2017) highlighted similar reductions. The resistant cultivar acted as an obstacle to spore dispersal in mixtures and thus induced a barrier effect on the susceptible one (Wolfe, 1985). This ‘protective’ effect has been reported before (Gigot *et al*., 2013) and was particularly strong in our study. Conversely, the presence of susceptible plants decreased the number of virulent strains filtered out by resistant plants relative to a pure stand of the resistant cultivar (Figure S11b and S11c). The susceptible plants did not concentrate more virulent strains in mixtures than in pure stands (Figure S11a and S11c), thereby decreasing the number of virulent strains at whole-canopy scale. This may explain the ‘protective’ effect of the susceptible cultivar on the resistant cultivar, which is a counter-intuitive but crucial result.

### Estimation of the local frequency of virulent strains

We estimated the local frequency of virulent strains (0.13 in 2018 and 0.14 in 2019) using the pathogen population collected on pure stands of the susceptible cultivar early in the epidemic, which should reflect the founding ascospore population (Figure 1b). Some of the strains collected from sporulating lesions on Cellule early in the epidemic were avirulent. Two hypotheses may explain this: (i) the ontogeny or temperature-modulation of *Stb16q* expression in field conditions, and (ii) the possibility of an avirulent strain co-infecting a host plant by taking advantage of infection with a virulent strain (‘stowaway effect’) or following systemic induced susceptibility (Seybold *et al*., 2020).

### Evolution of the frequency of virulent strains in mixtures

Mixtures modified the composition of the initial pathogen population, by affecting strain selection during the epidemic. The frequency of virulent strains collected on the susceptible cultivar did not differ between pure stands and mixtures (Figure 5) early in the epidemic, when secondary infections were not yet strongly affecting disease dynamics. The differences expressed later, after several cycles of secondary infection, suggest that the mixture effect results from cross-contaminations between cultivars driven by pycnidiospores splash dispersal (Figure 1d), consistent with previous findings (Gigot *et al*., 2013; Vidal *et al*., 2017). In this study, the optimal proportion of the resistant cultivar minimising the size of the virulent subpopulation 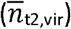 and the frequency of virulent strains 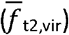 appeared to be about 0.25 (Figure 7e and f7). The overall effect would probably be different for mixtures of cultivars with different combinations of resistance genes. For determining the consequences of multiple virulence-resistance interactions, a parallel could be drawn with studies focusing on the impact of wheat mixtures on the structure of yellow rust and powdery mildew populations (Villaréal & Lannou, 2000; Lannou *et al*., 2005) and on the emergence of multivirulent strains. It was shown that the frequency of such strains was not necessarily significantly higher in mixtures than in pure stands (Chin & Wolfe, 1984; Dileone & Mundt, 1994).

### No evidence of a fitness cost associated with AvrStb16q virulence

In theory, there should be a ‘fitness penalty’ for pathogen populations with high levels of virulence after adaptation to hosts (Laine & Barrès, 2013). We did not find such a cost of virulence against *Stb16q* related to the aggressiveness trait that we studied (percentage of leaf area covered by pycnidia). This may be because the experimental conditions resulted in small mean sporulating areas (not entirely suitable for such a test), because this virulence has no cost — consistently with the experimental findings of Cowger & Mundt, 2002b — or because the cost concerns another aggressiveness trait (e.g. latent period, sporulation capacity of a pycnidia, infection efficacy; Suffert *et al*., 2013). Further studies, specially designed for this purpose, would help to progress on this issue.

### Impact of the proportion of cultivars on mixture effects

We showed that changes on the overall size and composition of the pathogen population depended on the proportions of the two cultivars, consistent with previous findings (Finckh *et al*., 2000). All the mixtures efficiently decreased the size of the pathogen population 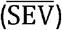, but the decrease in severity relative to expected levels in the absence of a mixture effect (Figure 4) was greatest for C1A1 in both 2018 (40%) and 2019 (71%). In our experiment, the optimal proportion of the resistant cultivar for reducing 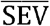 seemed to lie between 0.5 and 0.75. This is consistent with simulations performed by Gigot *et al*. (2014), which showed the ‘protection’ of susceptible plants by resistant plants to be optimal for mixtures with more resistant than susceptible plants.

### Impact of the intensity of the epidemic on mixture effects

The intensity of epidemics, driven by climatic conditions, may account for the observed differences in the magnitude of the mixture effect. The epidemic started earlier in 2018 (Figure 4) and this year was wetter and cooler than 2019. However, STB severity ended up being more intense on flag leaves in 2019 (19% in 2018 vs. 33% in 2019 for SEV_A_ in C0A1; data not shown). This apparent contradiction can be explained by late spring rainfall at key moments for the epidemic, while the primary inoculum is rarely a limiting factor (Morais *et al*., 2016). Mixtures decreased the size of the total pathogen population more effectively in 2019. Mixtures also had a greater impact on the frequency of virulent strains on the susceptible cultivar relative to the pure stand (Figure 5) in 2018 than in 2019, whereas the opposite pattern might have been expected, due to more intense disease dynamics late in the epidemic (Mundt, 2002). Gigot *et al*. (2013) showed that mixtures significantly decreased disease severity in most cases, regardless of disease intensity, whereas Kristoffersen *et al*. (2019) found no correlation with mixture efficacy.

### A trade-off between disease control and resistance durability?

The choice of the indicator to analyse and the pure stand of reference is crucial. Our findings suggest that the optimal proportions of cultivars for reducing the size of the virulent subpopulation 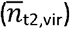 may differ slightly from those for reducing the size of the total pathogen population 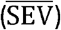, and those for reducing the overall frequency of virulent strains 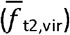 (Figure 7). The potential differences in the optima for these three indicators may conceal a trade-off between disease control and resistance durability. When comparing mixtures to the pure stands of the susceptible cultivar, disease severity can, indeed, be reduced by including a resistant cultivar in the host population (Figure 4). However, this effect may be counterbalanced by an acceleration of the breakdown dynamics of this resistance, due to an increase in the frequency of virulent strains in susceptible plants. The conclusion would clearly be different if a pure stand of the resistant cultivar is used as the reference: the inclusion of susceptible plants decreased the selection pressure exerted by emerging virulent strains, but also, as shown here, decreased disease severity on the resistant plants. The pathogen population on the susceptible plants of a cultivar mixture may be smaller than that in a pure stand or in a mixture with a high proportion of the susceptible cultivar, without there necessarily being an impact on the number of virulent strains (Figure S11). From this perspective, mixtures appeared to slow down the breakdown dynamics of the resistance. Moreover, we did not find a trade-off between the size of the virulent subpopulation and the frequency of virulent strains, but there is no reason why this trend should be the same in other contexts. The ‘size’ vs. ‘frequency’ debate is a classical issue in ecology that should not be neglected here, with consequences for genetic diversity and evolution (Brown & Tellier, 2011). Further studies are needed to determine if these dynamics would be different depending on the local frequency of virulent strains, i.e. the ‘breakdown stage’ of the resistance.

#### Perspectives

Determination of the effects of mixtures on the durability of the resistances deployed requires a consideration of the complete disease cycle, including sexual reproduction, and an extension of this approach to time scales beyond a single annual epidemic. One of the major issues arising from this study and previous studies (e.g., Finckh *et al*., 2000; Mundt, 2002; Lannou *et al*., 2005) is whether a cropping system based on cultivar mixtures would be likely to increase the prevalence of virulent strains after several years. This is both a question of the ‘reference’ used (pure stand of a susceptible or resistant cultivar) and of the objective assigned to the mixture: decreasing disease severity relative to the pure stand of susceptible plants or increasing the durability of resistance by reducing the pressure exerted on the virulent subpopulation of the pathogen. Investigations of the longer-term effects of cultivar mixtures need to consider sexual reproduction, as this process may either amplify or attenuate the demogenetic changes occurring during an annual epidemic. This is all the more relevant because a trade-off between size of pathogen population and frequency of virulence would impact the efficacy and durability, respectively, of the cultivar deployment strategies.

Recent landscape-scale models have investigated the effect of the proportion of fields sown with resistant and susceptible cultivars and their connectivity on the dynamics of resistance breakdown after several generations (e.g. Fabre *et al*., 2015). In comparisons of different strategies for preventing or delaying the appearance of a ‘superpathogen’, cultivar mixtures appeared to be effective in the long term for pathogens with high mutation rates and high fitness costs (Rimbaud *et al*., 2018; Crété *et al*., 2020). Our experimental findings confirm that cultivar mixtures may also be used to extend the lifetime of resistance sources, this time at within-field scale, through the introduction of heterogeneity.

## Acknowledgements

We thank Christophe Montagnier (INRAE Experimental Unit, Thiverval-Grignon, FR), Nathalie Retout, Laurent Gerard, Nicolas Lecutier, Auriane Pinton, Miléna Bonnard, Jérôme Lageyre, Martin Willigsecker and Benjamin Boudier (INRAE BIOGER, Thiverval-Grignon, FR) for technical assistance. We thank Dr. Cyrille Saintenac (INRAE GDEC, Clermont-Ferrand, FR) for providing us with the Chinese Spring isogenic lines of wheat. We thank Dr. Romain Valade (ARVALIS-Institut du Végétal, Thiverval-Grignon, FR) and Dr. Thierry Marcel (INRAE BIOGER, Thiverval-Grignon, FR) for sharing information and providing relevant advices throughout this project. We thank Dr. Julie Sappa for her help correcting our English.

## Data Availability Statement

The data and the R scripts that support the findings of this study [Files F1-21, see list below] are freely available from the INRAE Dataverse online data repository (https://data.inrae.fr/) at https://doi.org/10.15454/4MAAI0.

**File F1 *(F1_Field_disease_severity_rawdata*.*csv)***. Field disease severity raw data.

**File F2 *(F2_t2_2018_field-severity_means*.*txt)***. Disease severity means per block, treatment, and cultivar used to calculate a proxy for the size of the virulent subpopulations. Data for the late stage of 2018 epidemic.

**File F3 *(F3_t2_2019_field-severity_means*.*txt)***. Disease severity means per block, treatment, and cultivar used to calculate a proxy for the size of the virulent subpopulations. Data for the late stage of 2019 epidemic.

**File F4 *(F4_List_of_isolates*.*xls)***. List and virulence status of the 3035 isolates phenotyped in the greenhouse.

**File F5 *(F5_Disease_score_population-phenotyping_rawdata*.*xls)***. Disease score raw data for the population-phenotyping.

**File F6 *(F6_t2_2018_population-phenotyping_virulents_number_R_on_R*.*txt)***. Contingency table with the number of virulent and avirulent isolates per block and treatment, collected on Cellule and tested on Cellule, characterized with the population-phenotyping. Data for the late stage of 2018 epidemic.

**File F7 *(F7_t2_2018_population-phenotyping_virulents_number_R_on_S*.*txt)***. Contingency table with the number of pathogenic and non-pathogenic isolates per block and treatment, collected on Cellule and tested on Apache, characterized with the population-phenotyping. Data for the late stage of 2018 epidemic.

**File F8 *(F8_t2_2018_population-phenotyping_virulents_number_S_on_R*.*txt)***. Contingency table with the number of virulent and avirulent isolates per block and treatment, collected on Apache and tested on Cellule, characterized with the population-phenotyping. Data for the late stage of 2018 epidemic.

**File F9 *(F9_t2_2018_population-phenotyping_virulents_number_S_on_S*.*txt)***. Contingency table with the number of pathogenic and non-pathogenic isolates per block and treatment, collected on Apache and tested on Apache, characterized with the population-phenotyping. Data for the late stage of 2018 epidemic.

**File F10 *(F10_t2_2018_TabContglobal_virulence*.*txt)***. Global contingency table with the number of virulent, avirulent, pathogenic and non-pathogenic isolates per block, treatment, and cultivar collected on field at the late stage of 2018 epidemic and tested with the population-phenotyping. Data is used to calculate a proxy for the size of the virulent subpopulations.

**File F11 *(F11_t2_2019_population-phenotyping_virulents_number_R_on_R*.*txt)***. Contingency table with the number of virulent and avirulent isolates per block and treatment, collected on Cellule and tested on Cellule, characterized with the population-phenotyping. Data for the late stage of 2019 epidemic.

**File F12 *(F12_t2_2019_population-phenotyping_virulents_number_R_on_S*.*txt)***. Contingency table with the number of pathogenic and non-pathogenic isolates per block and treatment, collected on Cellule and tested on Apache, characterized with the population-phenotyping. Data for the late stage of 2019 epidemic.

**File F13 *(F13_t2_2019_population-phenotyping_virulents_number_S_on_R*.*txt)***. Contingency table with the number of virulent and avirulent isolates per block and treatment, collected on Apache and tested on Cellule, characterized with the population-phenotyping. Data for the late stage of 2019 epidemic.

**File F14 *(F14_t2_2019_population-phenotyping_virulents_number_S_on_S*.*txt)***. Contingency table with the number of pathogenic and non-pathogenic isolates per block and treatment, collected on Apache and tested on Apache, characterized with the population-phenotyping. Data for the late stage of 2019 epidemic.

**File F15 *(F15_t2_2019_TabContglobal_virulence*.*txt)***. Global contingency table with the number of virulent, avirulent, pathogenic and non-pathogenic isolates per block, treatment, and cultivar collected on field at the late stage of 2019 epidemic and tested with the population-phenotyping. Data is used to calculate a proxy for the size of the virulent subpopulations.

**File F16 *(F16_Severity_analysis*.*R)***. Main script for field data analysis.

**File F17 *(F17_Severity_function*.*R)***. Function used in main script for field data analysis.

**File F18 *(F18_Virulence_analysis*.*R)***. Main script for population-phenotyping data analysis.

**File F19 *(F19_Virulence_function*.*R)***. Function used in main script for population-phenotyping data analysis.

**File F20 *(F20_Realpop_analysis*.*R)***. Main script to calculate the proxy for the size of the virulent subpopulations.

**File F21 *(F21_Realpop_function*.*R)***. Function used in main script to calculate the proxy for the size of the virulent subpopulations.

## Supplementary Material

Supplementary material is available in the online version of this article.

## References

Borg J, Kiaer LP, Lecarpentier C, Goldrienger I, Saint-Jean S, Enjalbert S, 2018. Unfolding the potential of wheat cultivar mixtures: A meta-analysis perspective and identification of knowledge gaps. Field Crops Research 221, 298–313.

Brown JKM, Chartrain L, Lasserre-Zuber P, Saintenac C, 2015. Genetics of resistance to Zymoseptoria tritici and applications to wheat breeding. Fungal Genetics and Biology 79, 33–41.

Brown JKM, Tellier A, 2011. Plant-parasite coevolution: bridging the gap between genetics and ecology. Annual Review of Phytopathology 29, 345–367.

Chartrain L, Brading PA, Brown JKM, 2005. Presence of the Stb6 gene for resistance to septoria tritici blotch (Mycosphaerella graminicola) in cultivars used in wheat-breeding programmes worldwide. Plant Pathology 54, 134–143.

Chin K, Wolfe M, 1984. Selection on Erysiphe graminis in pure and mixed stands of barley. Plant Pathology 33, 535–545.

Cowger C, Hoffer ME, Mundt CC, 2000. Specific adaptation by Mycosphaerella graminicola to a resistant wheat cultivar. Plant Pathology 49, 445–451.

Cowger C, Mundt CC, 2002a. Effects of wheat cultivar mixtures on epidemic progression of Septoria tritici blotch and pathogenicity of Mycosphaerella graminicola. Phytopathology 92, 617–623.

Cowger C, Mundt CC, 2002b. Aggressiveness of Mycosphaerella graminicola isolates from susceptible and partially resistant wheat cultivars. Phytopathology 92, 624–630.

Crété R, Pires RN, Barbetti MJ, Renton M, 2020. Rotating and stacking genes can improve crop resistance durability while potentially selecting highly virulent pathogen strains. Scientific Reports 10, 19752.

Dai J, Wiersma JJ, Nolen DL, 2012. Performance of hard red spring wheat cultivar mixtures. Agronomy Journal 104, 17–21.

Dileone J, Mundt C, 1994. Effect of wheat cultivar mixtures on populations of Puccinia striiformis races. Plant Pathology 43, 917–930.

Fabre F, Rousseau E, Mailleret L, Moury B, 2015. Epidemiological and evolutionary management of plant resistance: optimizing the deployment of cultivar mixtures in time and space in agricultural landscapes. Evolutionary Applications 8, 919–932.

Finckh MR, 2008. Integration of breeding and technology into diversification strategies for disease control in modern agriculture. European Journal of Plant Pathology 121, 399– 409.

Finckh M, Gacek E, Goyeau H et al., 2000. Cereal variety and species mixtures in practice, with emphasis on disease resistance. Agronomie 20, 813–837.

Fones H, Gurr S, 2015. The impact of Septoria tritici blotch disease on wheat: An EU perspective. Fungal Genetics and Biology 79, 3–7.

Ghaffary SMT, Faris JD, Friesen TL et al., 2012. New broad-spectrum resistance to Septoria tritici blotch derived from synthetic hexaploid wheat. Theoretical and Applied Genetics 124, 125–142.

Gigot C, Saint-Jean S, Huber L et al., 2013. Protective effects of a wheat cultivar mixture against splash-dispersed Septoria tritici blotch epidemics. Plant Pathology 62, 1011– 1019.

Gigot C, de Vallavieille-Pope C, Huber L, Saint-Jean S, 2014. Using virtual 3-D plant architecture to assess fungal pathogen splash dispersal in heterogeneous canopies: a case study with cultivar mixtures and a non-specialized disease causal agent. Annals of Botany 114, 863–876.

Johnson R, 1984. A critical analysis of durable resistance. Annual Review of Phytopathology 22, 309–330.

Kildea S, Byrne JJ, Cucak M, Hutton F, 2020. First report of virulence to the Septoria tritici blotch resistance gene Stb16q in the Irish Zymoseptoria tritici population. New Disease Reports 41, 13–13.

Kristoffersen R, Jørgensen LN, Eriksen LB, Nielsen GC, Kiær LP, 2019. Control of Septoria tritici blotch by winter wheat cultivar mixtures: Meta-analysis of 19 years of cultivar trials. bioRxiv, 658575.

Laine A-L, Barrès B, 2013. Epidemiological and evolutionary consequences of life-history trade-offs in pathogens. Plant Pathology 62, 96–105.

Lannou C, 2012. Variation and selection of quantitative traits in plant pathogens. Phytopathology 50, 319–38.

Lannou C, Hubert P, Gimeno C, 2005. Competition and interactions among stripe rust pathotypes in wheat-cultivar mixtures. Plant Pathology 54, 699–712.

McDonald BA, Linde C, 2002. Pathogen population genetics, evolutionary potential, and durable resistance. Annual Review of Phytopathology 40, 349–379.

McIntosh RA, Brown GN, 1997. Anticipatory breeding for resistance to rust diseases in wheat. Annual Review of Phytopathology 35, 311–326.

McRoberts N, Hughes G, Madden LV, 2003. The theoretical basis and practical application of relationships between different disease intensity measurements in plants. Annals of Applied Biology 142, 191–211.

Morais D, Sache I, Suffert F, Laval V, 2016. Is the onset of Septoria tritici blotch epidemics related to the local pool of ascospores? Plant Pathology 65, 250–260.

Mundt CC, 2002. Use of multiline cultivars and cultivar mixtures for disease management. Annual Review of Phytopathology 40, 381–410.

Niks RE, Parlevliet JE, Lindhout P, Bai Y, 2011. Breeding crops with resistance to diseases and pests. Wageningen: Wageningen Academic Publishers.

Perronne R, Makowski D, Goffaux R, Montalent P, Goldringer I, 2017. Temporal evolution of varietal, spatial and genetic diversity of bread wheat between 1980 and 2006 strongly depends upon agricultural regions in France. Agriculture, Ecosystems & Environment 236, 12–20.

Rimbaud L, Papaix J, Barrett LG, Burdon JJ, Thrall PH, 2018. Mosaics, mixtures, rotations or pyramiding: What is the optimal strategy to deploy major gene resistance? Evolutionary Applications 11, 1791–1810.

Seybold H, Demetrowitsch TJ, Hassani MA et al., 2020. A fungal pathogen induces systemic susceptibility and systemic shifts in wheat metabolome and microbiome composition. Nature Communications 11, 1–12.

Suffert F, Delestre G, Gélisse S, 2018. Sexual reproduction in the fungal foliar pathogen Zymoseptoria tritici is driven by antagonistic density dependence mechanisms. Microbial Ecology 77, 110–123.

Suffert F, Sache I, Lannou C, 2011. Early stages of Septoria tritici blotch epidemics of winter wheat: build-up, overseasoning, and release of primary inoculum. Plant Pathology 60, 166–177.

Suffert F, Sache I, Lannou C, 2013. Assessment of quantitative traits of aggressiveness in Mycosphaerella graminicola on adult wheat plants. Plant Pathology 62, 1330–1341.

Vidal T, Boixel A-L, Durand B, de Vallavieille-Pope C, Huber L, Saint-Jean S, 2017. Reduction of fungal disease spread in cultivar mixtures: Impact of canopy architecture on rain-splash dispersal and on crop microclimate. Agricultural and Forest Meteorology 246, 154–161.

Villaréal LMMA, Lannou C, 2000. Selection for increased spore efficacy by host genetic background in a wheat powdery mildew population. Phytopathology 90, 1300–1306.

Wolfe MS, 1985. The current status and prospects of multiline cultivars and variety mixtures for disease resistance. Annual Review of Phytopathology 23, 251–273.

