## Supplementary material for "Annual dynamics of *Zymoseptoria tritici* populations in wheat cultivar mixtures: a compromise between the efficiency and durability of a recently broken-down resistance gene?"

**Table S1.** Actual proportion of wheat plants of cv. Apache (A) and cv. Cellule (C) in microplots in pure stands (C0A1 and C1A0) and in mixtures (C1A3, C1A1, and C3A1) at the late stage of epidemics in 2018 and 2019. Each proportion was estimated from the proportion of wheat spikes of each cultivar at heading assessed on the basis of the presence (C) or absence (A) of bearded heads (counted along three 1-m lines on July 4^th^, 2018 and July 1^st^, 2019). The total number of spikes per metre is indicated in italics, in brackets.

| Year | Block | C0A1 | C1A3 | | | C1A1 | | | C3A1 | | | | | C1A0 |
| --- | --- | --- | --- | --- | --- | --- | --- | --- | --- | --- | --- | --- | --- | --- |
|  |  | A | C | | A | C | A | | C | A | | | | C |
| 2018 | a | 1 | 0.12 | | 0.88 | 0.52 | 0.48 | | 0.61 | 0.39 | | 1 | | |
|  |  | *(82)* | *(68)* | | | *(68)* | | | *(75)* | | | *(79)* | | |
|  | b | 1 | 0.25 | | 0.75 | 0.27 | 0.73 | | 0.71 | 0.29 | | 1 | | |
|  |  | *(76)* | *(82)* | | | *(75)* | | | *(92)* | | | *(82)* | | |
|  | c | 1 | 0.20 | | 0.80 | 0.46 | 0.54 | | 0.69 | 0.31 | | 1 | | |
|  |  | *(67)* | *(82)* | | | *(77)* | | | *(73)* | | | *(97)* | | |
|  | mean | 1 | 0.19 | 0.81 | | 0.42 | | 0.58 | 0.67 | | 0.33 | | 1 | |
|  |  | *(75)* | *(77)* | | | *(73)* | | | *(80)* | | | *(86)* | | |
| 2019 | a | 1 | 0.14 | | 0.86 | 0.35 | 0.65 | | 0.76 | 0.24 | | 1 | | |
|  |  | *(65)* | *(86)* | | | *(69)* | | | *(74)* | | | *(79)* | | |
|  | b | 1 | 0.39 | | 0.61 | 0.60 | 0.40 | | 0.61 | 0.39 | | 1 | | |
|  |  | *(103)* | *(77)* | | | *(82)* | | | *(95)* | | | *(73)* | | |
|  | c | 1 | 0.31 | | 0.69 | 0.35 | 0.65 | | 0.82 | 0.18 | | 1 | | |
|  |  | *(91)* | *(72)* | | | *(65)* | | | *(95)* | | | *(69)* | | |
|  | mean | 1 | 0.28 | 0.72 | | 0.43 | | 0.57 | 0.73 | | 0.27 | | 1 | |
|  |  | *(86)* | *(78)* | | | *(72)* | | | *(88)* | | | *(73)* | | |

**Table S2.** The origin (treatment, from the following list: C1A3, C1A1, C3A1, C1A0; period, t1 or t2, in 2018) of the 47 *Z. tritici* strains shown to be avirulent against *Stb16q* established in the first population-phenotyping, for which virulence status was validated by confirmatory individual phenotyping. The total number of strains tested is indicated in brackets. There was no significant difference in the number of avirulent strains at t1 between treatments (Fisher’s test, *p* = 1).

|  | C1A3 | C1A1 | C3A1 | C1A0 |
| --- | --- | --- | --- | --- |
| t1 | 7 (11) | 7 (11) | 4 (7) | 6 (12) |
| t2 | 0 (4) | 0 (1) | 0 (0) | 0 (1) |

**Table S3.** Main sources of uncertainty and benefit-disadvantage analysis of the ‘population phenotyping’ method compared to the classical ‘confirmatory individual phenotyping’ method (see also Figure S3). ‘False negative’ refers to virulent strains that appeared avirulent in the population phenotyping but were actually virulent in the confirmatory individual phenotyping.

| Sources of uncertainty | Disadvantages | Solutions for reducing disadvantages and estimating uncertainty | Benefits |
| --- | --- | --- | --- |
| Preparation of the inoculum suspensions | The targeted concentration 10^5^ spores·mL^-1^ is neither precisely measured nor adjusted, resulting in a loss of accuracy. | The mean concentration of suspensions was estimated ±3.0 10^5^ spores·mL^-1^ from a sample of 40 strains, with 85% in the range 1.0 x 10^5^-5.0 x 10^5^ spores·mL^-1^ (Figure S4). The impact of such variation (see Figure 8 in Suffert *et al.*, 2018) can be considered negligible for characterising the virulence status of strains and acceptable for comparing the orders of magnitude of aggressiveness between virulent and avirulent strains. Uncertainty was further reduced by having the same person prepare all the batches of inoculum suspensions. | Overall preparation time cut to a third (±30 isolates vs. >100 isolates per half day) |
| Climatic conditions in the greenhouse | Semi-controlled environmental conditions (temperature, moisture and light) can only be maintained within a range of variation and not set at given values. The outdoor climatic conditions may influence the climatic conditions in the greenhouse and, thus, the physiology of wheat plants between batches due to seasonality. | The impact of climatic conditions on the development of *Z. tritici* *in planta* has been clearly established (McCorison & Goodwin, 2020). Hygrometric variations and variations of light exposure may explain some of the ‘false negatives’ and the poor results obtained with two non-conforming batches. Comparing these less suitable conditions with a classical phenotyping method in a growth chamber, a low threshold was used to identify strains as virulent (at least 5% of the area sporulating on at least one of the six inoculated leaves). Air temperature monitoring (Figure S5) made it possible to check that the thermal time from inoculation to disease assessment was significantly longer than the mean latent period of *Z. tritici* for all batches. The two non-conforming batches were identified (using eight virulent strains as control), and removed before retesting. | Overall time for *in planta* phenotyping reduced by at least 40% due to space saving in our facilities |
| Small number of biological replicates:  - Only six leaves inoculated  - Potential leaf layer effect  - No repetition of the batches | The number of ‘false negatives’ is expected to be inversely related to the number of biological replicates (number of leaves inoculated with a given strain) and to be higher in the absence of repetitions. | Confirmatory individual phenotyping was performed with a larger number of biological replicates in more suitable conditions (growth chamber). Half the strains collected on cv. Cellule in the field that appeared avirulent after the first population-phenotyping were actually virulent. Thus, the maximal percentage of ‘false negatives’ was estimated ±3%, which was considered acceptable with respect to the goal of this study. | Overall time for *in planta* phenotyping reduced by at least 40% due to space saving in our facilities.  Possibility of testing a very large number of strains (several thousand rather than several hundred) |
| Visual assessment | Visual assessment is theoretically less accurate than image analysis for area estimation (e.g. Stewart *et al.*, 2016). | The difficulties detecting pycnidia made it impossible to use image analysis in practice for high-throughput population phenotyping on seedlings. Uncertainty was reduced further by having the same person perform all visual assessments. | Overall time for *in planta* phenotyping reduced by at least 40% in our facilities |

**Figure S1**. Design of the field experiment performed at the INRAE experimental station in Grignon (Yvelines, France, 48°51′N, 1°58′E) in 2018 and repeated in 2019. The field plot was previously under maize and ploughed. It was divided into three blocks (a, b, c), each consisting in five microplots (3.5 m wide x 8 m long, with an inter-row spacing of 0.175 m) corresponding to five randomly distributed treatments: a pure stand of wheat cv. Apache (C0A1; 220 Apache seeds·m^-2^), a pure stand of wheat cv. Cellule (C1A0; 220 Cellule seeds·m^-2^), and three binary mixtures with proportions of Cellule of 0.25 (C1A3; 55 Cellule seeds·m^-2^ + 165 Apache seeds·m^-2^), 0.5 (C1A1; 110 Cellule seeds·m^-2^ + 110 Apache seeds·m^-2^) and 0.75 (C3A1; 165 Cellule seeds·m^-2^ + 55 Apache seeds·m^-2^). Seed lots were prepared with the established proportions by taking into account the specific weight of each cultivar seed (TKW Apache = 45 g; TKW Cellule = 46 g). The microplots were separated by 5.25 m-wide strips of triticale cv. Vuka (220 seeds·m^-2^) resistant to *Z. tritici* to prevent disease spread between neighbouring plots. Each microplot was split longitudinally into two equal parts. In the first part (western side), lines 2-3-4 were used for strain collection, and lines 6-7-8 for the estimation of disease severity early in the epidemic (t1); in the second part (eastern side), lines 2-3-4 and lines 6-7-8 were used similarly at the late stage of the epidemic (t2), and lines 6-7-8 for the estimation of the actual proportions of each cultivar (spikes).

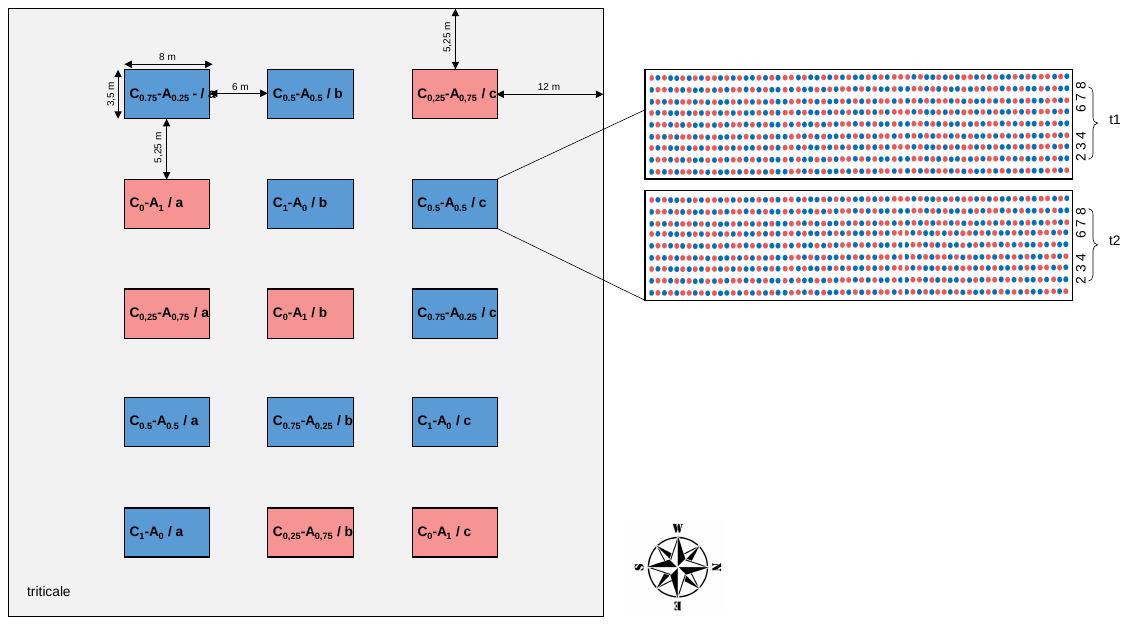

**Figure S2**. Overview of field conditions during disease assessments and sampling at the early (t1) and late (t2) stages of the epidemic. (a) Microplot at t1 (March 7, 2018). (b) Wheat seedlings of cv. Apache (red seed) and cv. Cellule (green seed). (c) Sporulating lesion caused by *Z. tritici* on an Apache seedling. (d) Microplot at t2 (May 11, 2018). (e) Adult plants of Apache (head not bearded) and Cellule (bearded head). (f) Cluster of lesions on an Apache adult leaf.

**Figure S3**. Simplified method of inoculation developed for ‘population-phenotyping’, estimated to be twice as efficient as the classical method (30 isolates per half day x 36 days vs. 100 isolates per half day x 22 days).

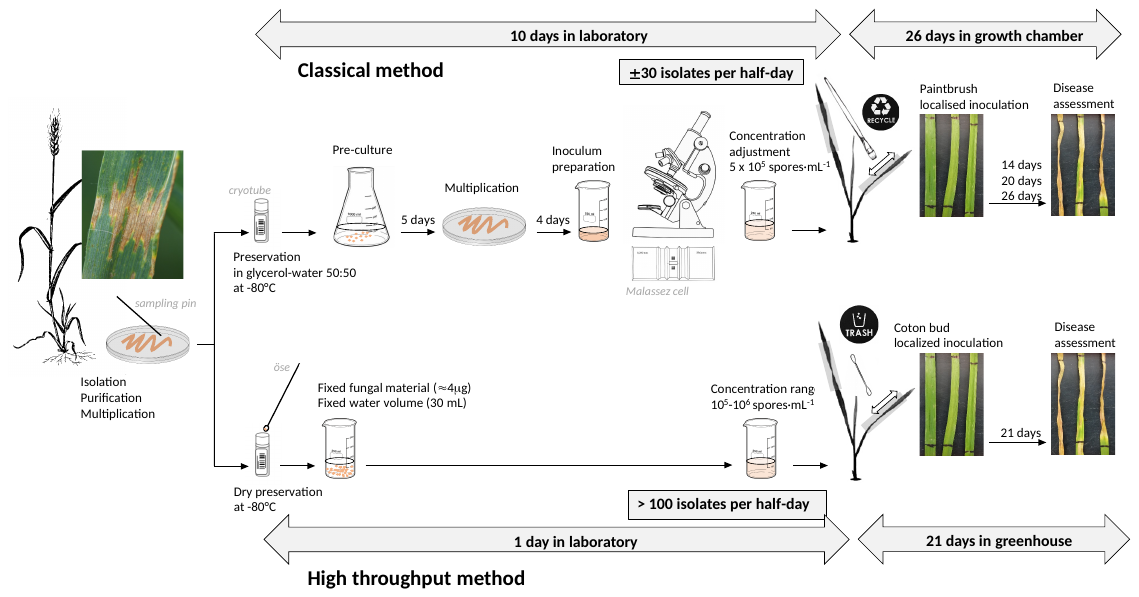

**Figure S4**. Variability of *Z. tritici* blastospore concentration in the non-adjusted suspensions used for ’population-phenotyping’. The simplified method involved diluting, in a volume of 30 mL of water, the mass of blastospores recovered with a 10-μL inoculating loop (central ring half-filled i.e., about 10 mg) from the freshly thawed contents of cryotubes stored at -80°C. Two independent series of tests (black and white dots) were performed with 2 x 20 strains. The mean concentration (white diamond) was 3.0 x 10^5^ spores·mL^-1^, with 85% of samples within the 10^5^-5.0 x 10^5^ spores·mL^-1^ range.

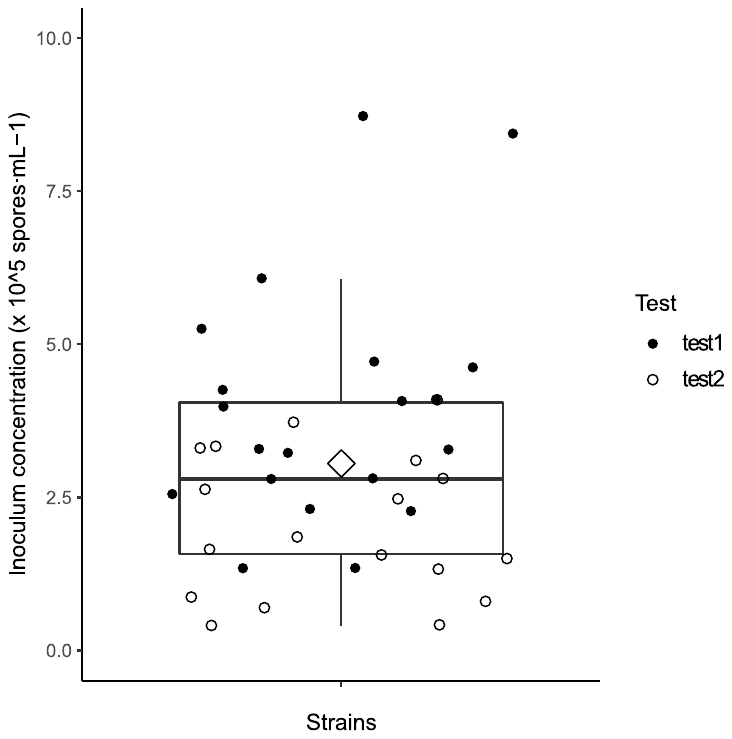

**Figure S5** Thermal conditions to which the 32 batches of *Z. tritici* strains were exposed during population-phenotyping in the greenhouse. Thermal time, expressed in degree-days post inoculation (°C·days^-1^), was calculated by summing daily mean air temperatures (0°C as the base temperature). The mean thermal time per population is plotted (white diamonds). Temperature was recorded every 15 min with a thermal sensor (digital probes with a SPY RF N recorder; JRI, Argenteuil, France). Batches were phenotyped (i) from March 12^th^ to June 6^th^, 2018 for strains collected at t1 in 2018, (ii) from June 25^th^ to August 21^st^ 2018 and from July 5^th^ to August 17^th^ 2019 for strains collected at t2 in 2018, (iii) from February 14^th^ to March 27^th^ and May 18^th^ to August 11^th^ 2020 for strains collected at t1 in 2019, (iv) from July 12^th^ to October 17^th^ 2019 for strains collected at t2 in 2019.

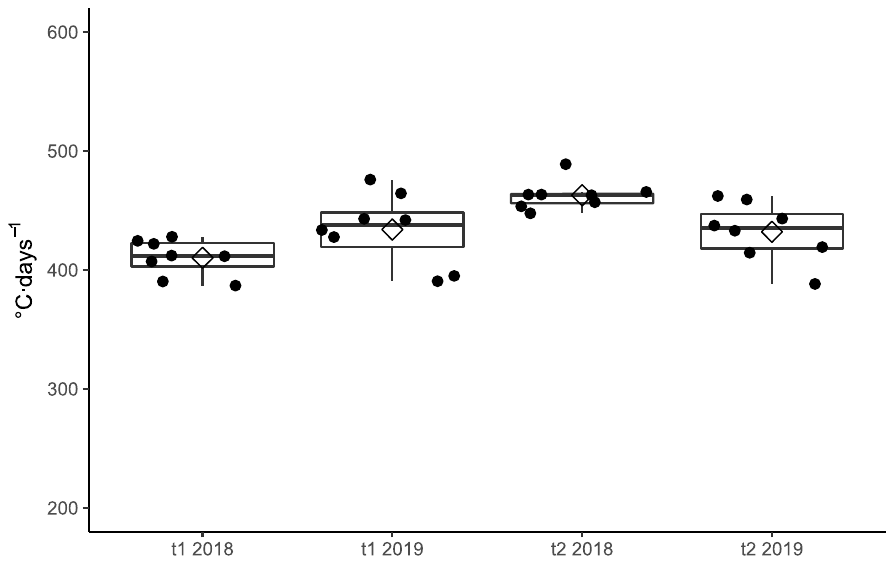

**Figure S6.** Frequency of *Z. tritici* strains considered ‘pathogenic’ (i.e., causing sporulating area on Apache; grey bar) and non-pathogenic (white bar) among the populations collected on Cellule in pure stands (p_C_ = 1) and mixtures (p_C_ = 0.25, 0.5 and 0.75 for C1A3, C1A1 and C3A1, respectively) at the early (t1) and late stages (t2) of the 2018 and 2019 epidemics. The number of pathogenic and non-pathogenic strains are indicated in grey and white boxes, respectively. The effect of treatment (pure stands and cultivar mixtures; letters at the top of each bar) on the number of pathogenic strains was assessed by performing Fisher’s exact test (the expected numbers were small), with Bonferroni correction for pairwise comparison.

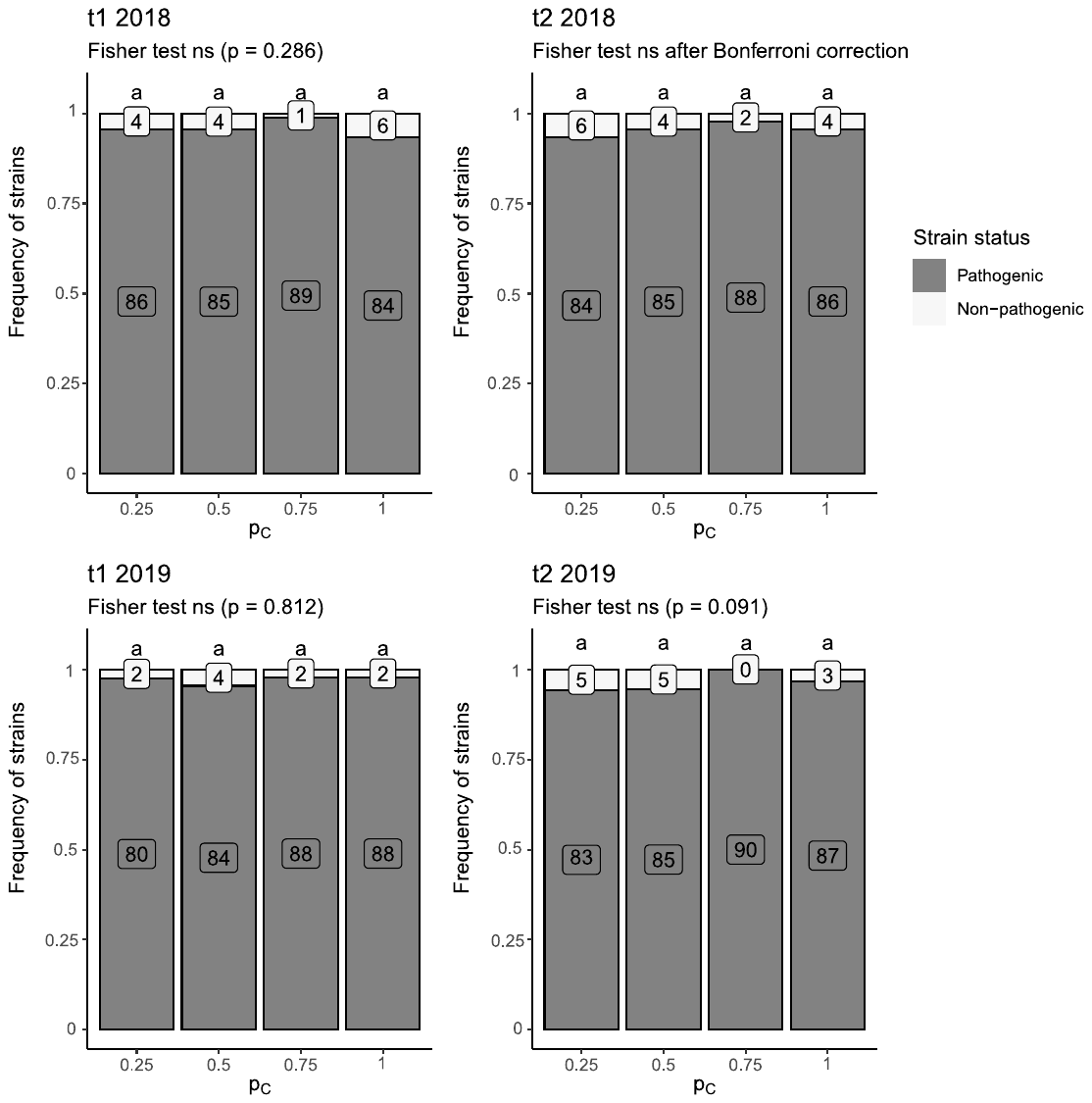

**Figure S7.** Frequency of *Z. tritici* strains considered pathogenic (i.e., causing sporulating area on Apache in the greenhouse; grey bars) and non-pathogenic (white bars) among the populations collected on Apache in pure stands (p_C_ = 0) and mixtures (p_C_ = 0.25, 0.50 and 0.75 for C1A3, C1A1 and C3A1, respectively) at the early (t1) and late stages (t2) of the 2018 and 2019 epidemics. The number of pathogenic and non-pathogenic strains are indicated in grey and white boxes, respectively. The effect of treatment (pure stands and cultivar mixtures; letters at the top of each bar) on the number of pathogenic strains was assessed by performing a χ² test, or Fisher’s exact test when the expected numbers were small, with Bonferroni correction for pairwise comparison.

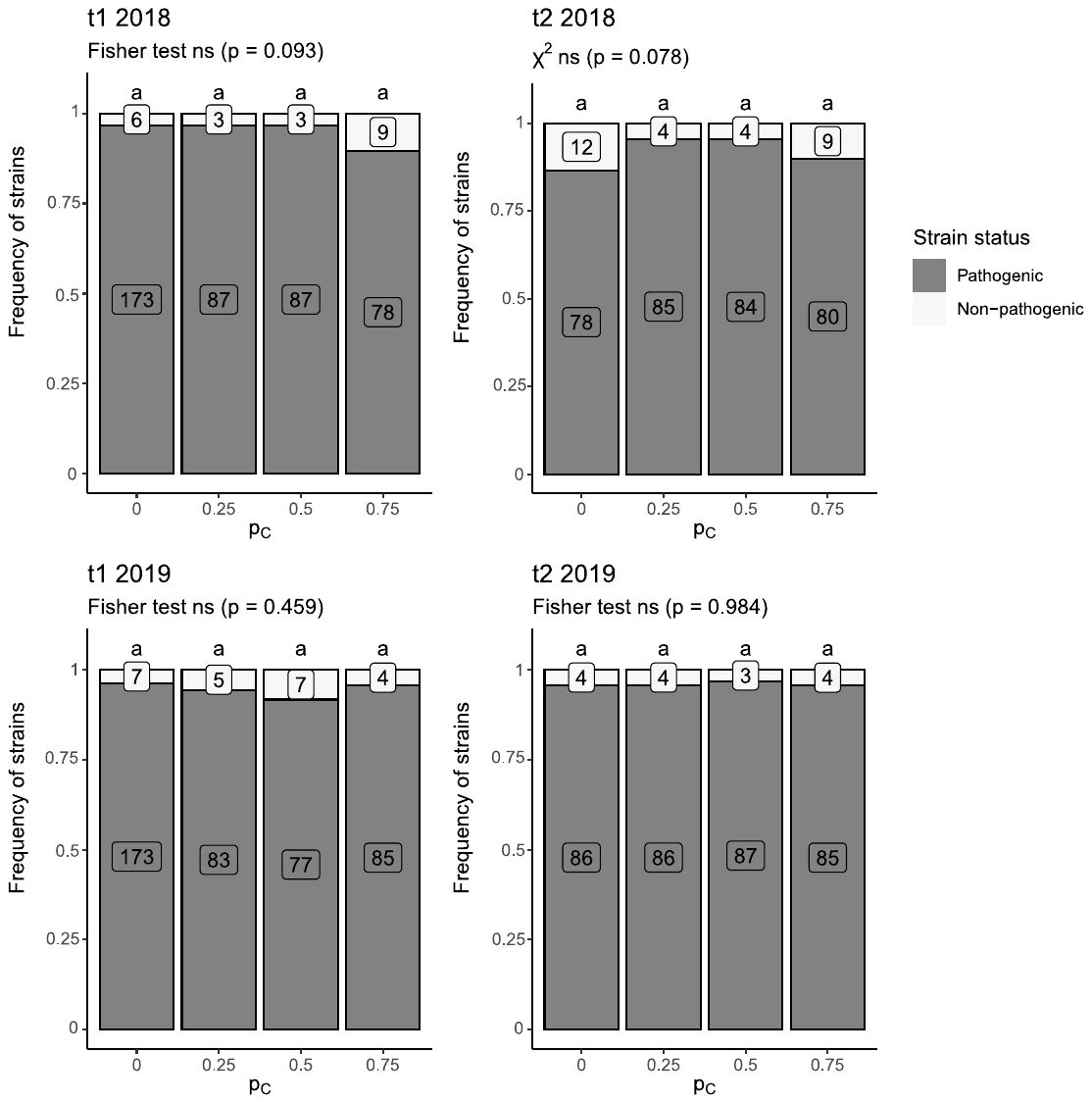

**Figure S8*.*** Comparison of the aggressiveness (AG for aggressiveness and AGc for conditional aggressiveness) of 2886 pathogenic strains according to their virulence status with respect to *Stb16q*. Aggressiveness, expressed as the percentage sporulating leaf area, was assessed on Apache. For AG, each dot represents the mean sporulating area induced by a strain on six leaves, and for AGc, the mean sporulating area on symptomatic leaves only. For each subpopulation the number of strains (n) and the mean are indicated. No significant difference between avirulent and virulent strains was observed for either AG (*p* = 0.054) or AGc (*p* = 0.060).

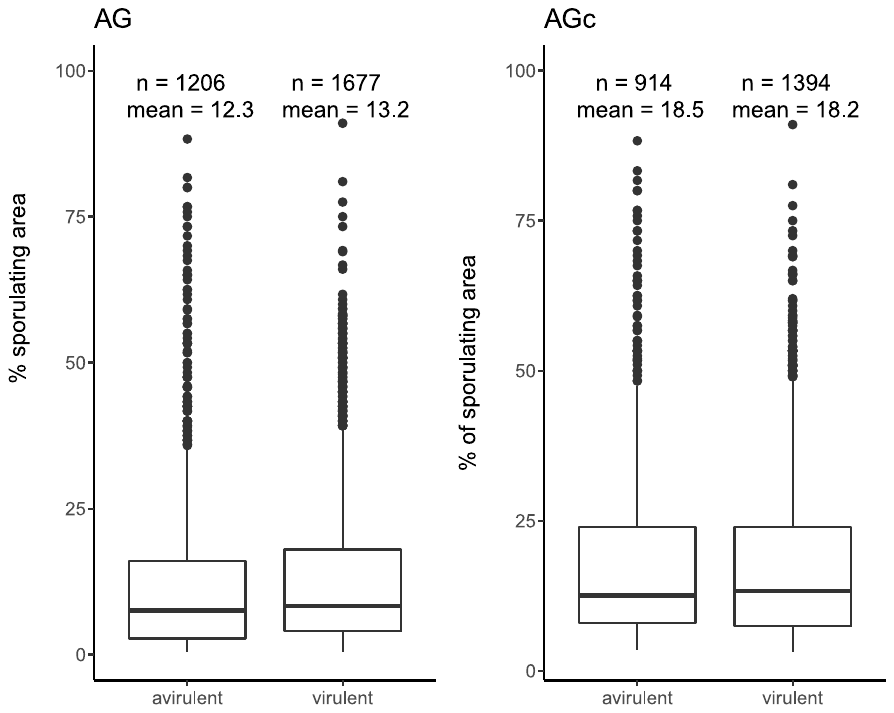

**Figure S9.** Aggressiveness of the 47 *Z. tritici* strains on Cellule (C) and on the isogenic line Chinese Spring with *Stb16q* (CS-*Stb16q*) assessed during confirmatory individual phenotyping. Each point is the mean percentage of the total area of the inoculated section covered with pycnidia on nine leaves per strain × wheat cultivar interaction. The virulence threshold was fixed at 10% (dashed lines). The 16 strains considered virulent on both Cellule and CS-*Stb16q* (mean aggressiveness 58.2% and 65.3%, respectively; Wilcoxon test, *p* < 0.0001) are indicated with black dots (yellow square). Two strains considered virulent only on Cellule and five strains considered virulent only on CS-*Stb16q* are indicated with white dots (blue rectangles). 24 strains considered avirulent on both Cellule and CS-*Stb16q* are indicated with crosses (red square).

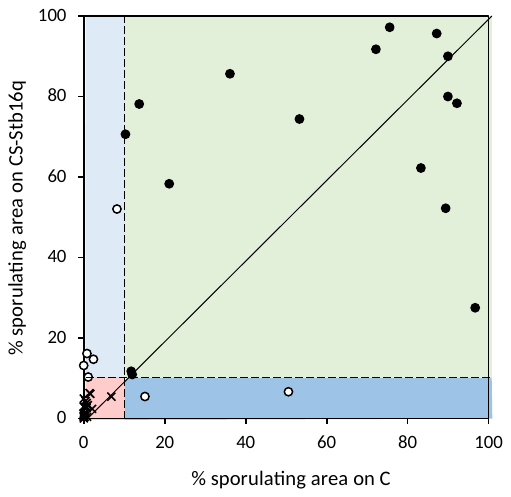

**Figure S10.** Analysis of disease severity data leading to the fixing at 10% of the relevant threshold (dashed line) for determining the virulence status of *Z. tritici* strains with respect to *Stb16q* 20 days post inoculation (dpi) in confirmatory individual phenotyping. Strains FS1766 (avirulent) and FS5948 (virulent) were tested three times (3 independent batches × 9 inoculated leaves as replicates) on the isolines Chinese Spring (CS), Chinese Spring with *Stb16q* (CS-*Stb16q*) and Cellule (C). Each point is the disease severity assessed on a leaf (i.e., sporulating area, expressed as a percentage of the inoculated leaf section).

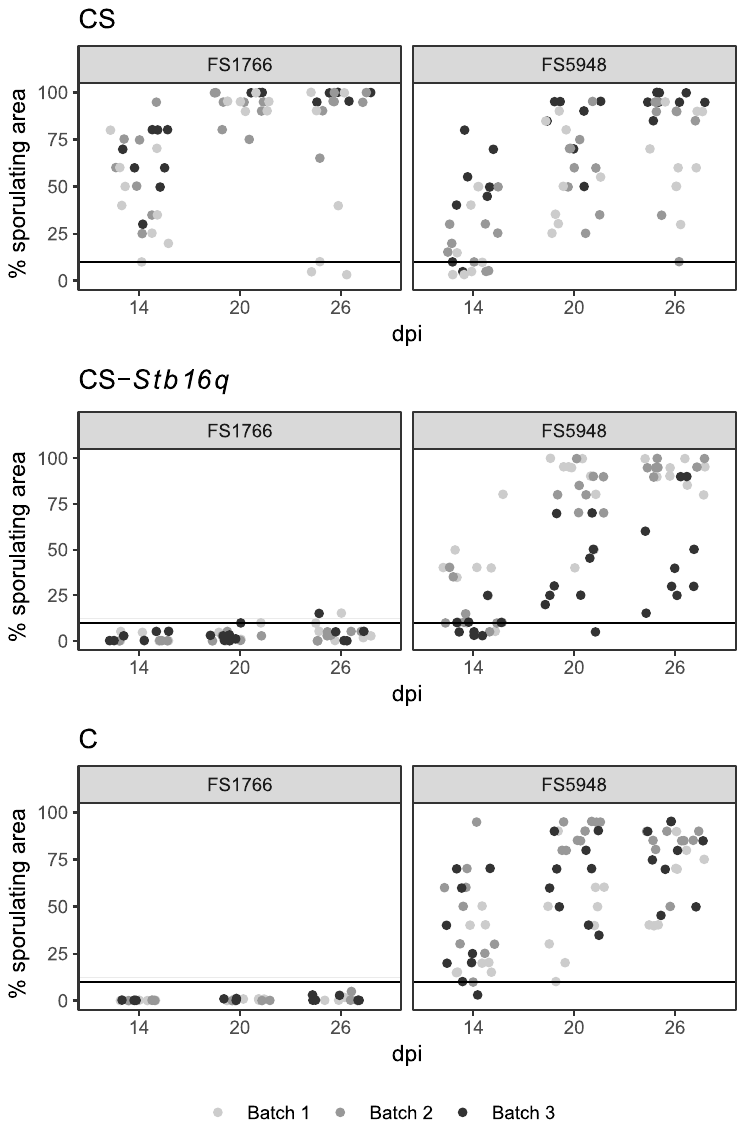

**Figure S11.** Impact of treatment (pure stands and mixtures) on the *Z. tritici* virulent subpopulation on each cultivar at the late stage of the epidemic (t2). (a, c) Mean size of the virulent subpopulation on Apache ($n$_t2,A,vir_; dimensionless variable) according to the proportion of Cellule (p_C_) in 2018 and 2019. (b, d) Mean size of the virulent subpopulation on Cellule ($n$_t2,C,vir_) according to the proportion of Cellule (p_C_) in 2018 and 2019. Grey dots indicate the mean value for each block and black dots indicate the mean for each treatment. Bars represent the standard error within each block.

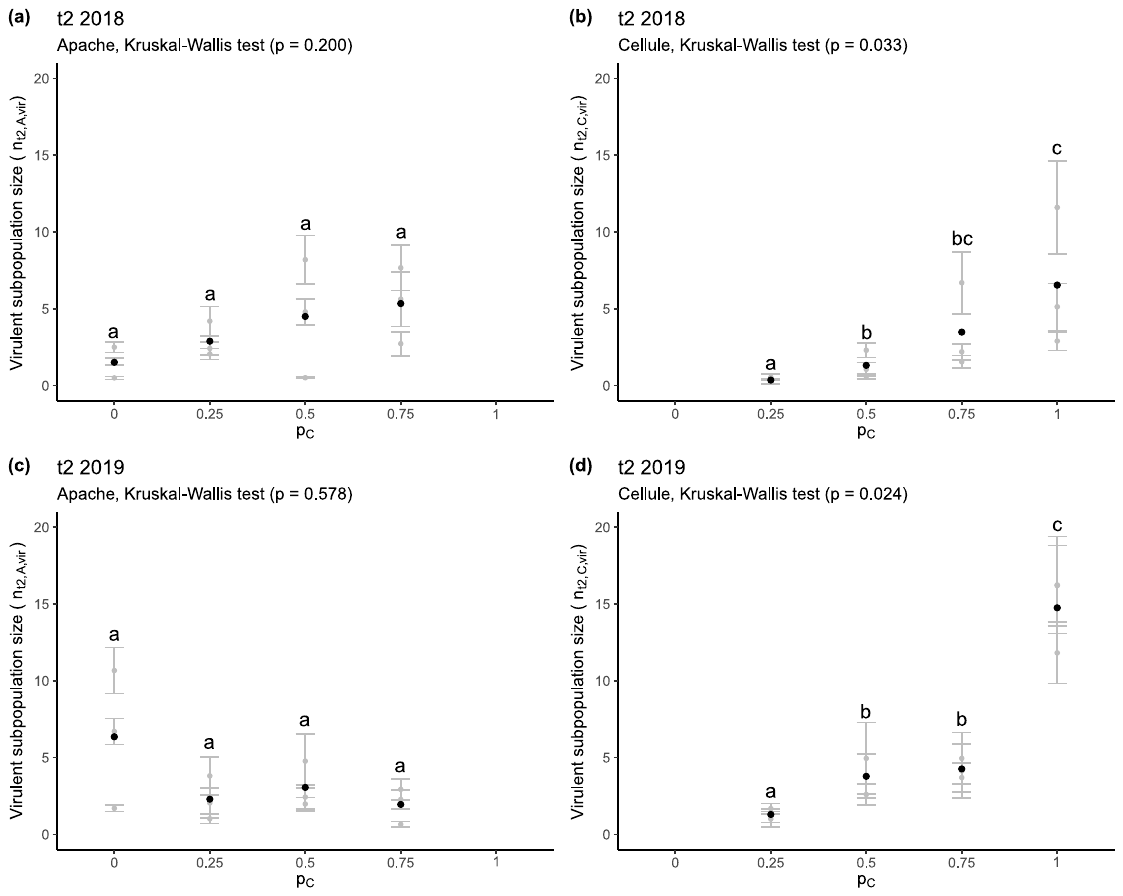
